# Estimation of Protein Melting Temperatures Using Small-Ladder Replica Exchange Simulations

**DOI:** 10.64898/2026.02.17.706302

**Authors:** Nithin K. Rajendran, Patrick K. Quoika, Martin Zacharias

## Abstract

The unfolding or melting temperature (*T*_*M*_) is a central quantity to characterize the stability of proteins and other biopolymers. The accurate prediction of protein melting temperatures by molecular mechanics force field simulations is highly desirable for many biophysical and biotechnological applications. Since the time scales for protein (un-)folding are hardly accessible in conventional MD (cMD) simulations, enhanced sampling techniques such as Temperature Replica Exchange Molecular Dynamics (TREMD) are typically employed. However, TREMD simulations are computationally very demanding especially if large temperature ranges need to be covered. Additionally, if the *T*_*M*_ is initially unknown, setting up TREMD simulations is often challenging. To find the optimal initial conditions for such simulations, we describe their performance based on a theoretical model, which we validate on a minimalistic Markov Chain Monte Carlo (MCMC) simulation setup. In an effort to reduce the computational demand, we have investigated the possibility to use small sets of TREMD temperature ladders placed iteratively in the vicinity of a *T*_*M*_ estimate. Different TREMD setups were extensively tested on the fast-folding protein Chignolin. We found that appropriate starting conformations lead to significantly faster convergence. Furthermore, we found that, in practice, combining multiple small temperature ladders can be advantageous in comparison to one single temperature ladder. Based on our findings, we formulate practical recommendations on how to set up TREMD for protein melting with optimal efficiency.

## 1 Introduction

Assessing the thermal stability of biomolecular secondary structures is important for several reasons. Practical considerations such as cold storage requirements, antibody solubility, and drug shelf-life are closely linked to thermal stability [1, 2]. At the same time, biomolecular function is tightly coupled to structure, while natural sequences are often optimized for activity rather than stability [3]. Characterizing thermal stability can therefore provide insights into the relationship between sequence, structure, and function, and can aid in the design of variants that demonstrate high stability and high activity. Similarly, systematic studies of stability changes induced by single-point mutations have been used to probe sequence–structure relationships in proteins [4].

Experimentally, thermal stability is commonly assessed using techniques such as circular dichroism (CD)[5] and differential scanning calorimetry (DSC) [6], differential scanning fluorimetry (DSF) [7], or other methods [8]. These methods typically involve monitoring temperature-dependent changes in some property such as the fraction of configurations or heat capacity. The melting temperature (*T*_*M*_) is identified as the mid-point of these transitions or as the point at which sharp transitions occur in the observed quantity. While reliable, these wet-lab methods are time-consuming and expensive for several reasons, including the need to prepare protein samples. Consequently, computational approaches are promising for complementing and (partially) replacing experimental efforts. The ease of introducing mutations or changes in environmental parameters in computational models enables high-throughput screening of biomolecular variants [9].

Atomistic Molecular Dynamics (MD) simulations have been widely employed to estimate *T*_*M*_ of biomolecules. However, several challenges limit their applicability. Simulation outcomes are sensitive to the choice of force field and water model [10, 11]. The time and length scale differences between real and simulated systems often prevents sufficient sampling of the free energy landscape. In the current age of exascale supercomputers, extensive probing of biomolecular systems can be performed in great detail. Time scales of the order of several tens of microseconds can be achieved on these machines, particularly when combined with AI/ML techniques [12]. However, for slow processes such as folding-unfolding, these time scales are still insufficient Mohr et al. [13].

Enhanced sampling methods in MD are routinely used to accelerate sampling of the phase space. Several enhanced sampling methods exist and their use depends on the quantity of interest. For example, simulated annealing is not suitable for estimating equilibrium properties such as melting temperatures, as the cooling rate influences final estimate. Other approaches such as replica exchange methods and metadynamics aim to improve sampling while preserving equilibrium statistics. The merits and demerits of these techniques have been extensively discussed in the literature [13, 14].

Temperature Replica Exchange Molecular Dynamics (TREMD) is a particularly popular approach for estimating melting temperatures of proteins and nucleic acids [15, 16]. TREMD achieves enhanced sampling by taking advantage of faster motion of atoms at higher temperatures. In TREMD, several copies of the system are simulated in parallel at increasing temperatures or at different “rungs” of a temperature “ladder”. At predetermined intervals, these copies are exchanged with each other using Metropolis Monte Carlo scheme. This allows the system to exploit accelerated kinetics at high temperatures while maintaining detailed balance. It has been suggested that the improvements in performance from TREMD would depend on the type of barrier to transitions; enthalpic barriers can be overcome more easily with TREMD than entropic ones [17]. However, TREMD scales poorly with system size and temperature range of interest. The method relies on frequent exchanges between neighboring replicas. Since the success rate of exchange attempts depends on the closeness of the potential energy distributions, the “rungs” cannot be placed too far apart.

In this study, we demonstrate that for estimating the melting temperatures of small biomolecules, it is not necessary to populate the entire temperature range of interest using a single, continuous temperature ladder. Instead, sets of disconnected temperature ladders containing as few as four to six replicas can be used. When combined with carefully chosen initial conditions, they can potentially provide reliable melting temperature estimates. While prior studies have examined the efficiency of TREMD relative to conventional MD (cMD), these analyses typically assume long simulation times and place little emphasis on the role of initialization [18–20]. Motivated in part by generative models that enable prediction of folded structures without extensive MD simulations, we investigate how the choice of starting structures can be optimized to accelerate convergence in small-ladder TREMD simulations. Our conclusions are supported by analytical arguments based on an Ornstein-Uhlenbeck (OU) model and validated computationally using TREMD and Markov Chain Monte Carlo (MCMC) simulations. Finally, the limitations of the approach are discussed in the context of novel systems.

## 2 Mathematical Model

The rate model of TREMD was developed by Rosta and Hummer [18] to compare TREMD simulations with cMD simulations. The efficiency of either method was quantified using the decay rate of the variance of an estimator of a state function. The specific case of systems dominated by slow transitions between two states was the primary focus. In their treatment, Rosta and Hummer [18] ignore the systematic effects of the choice of initial conditions. Consequently, the rate model forms an anchor for the present work, which aims to determine the effect of the set of starting structures on the convergence of TREMD simulations.

### 2.1 The Ornstein-Uhlenbeck Model

Consider a system that occupies metastable states folded (F) or unfolded (U). One might define the state using a range of values for some collective variable(s) such as the fraction of native contacts or root mean-squared distance with respect to a reference structure. For such a system, a discrete state function *s* can be defined such that

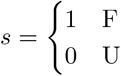

Using the collective variable and discrete state function, either 0 or 1 can be assigned to each frame in an MD trajectory at constant temperature *T* for a two-state system. The mean of this state function over a specified simulation time would be the estimate of the probability of occurrence of state F at temperature *T*. Over the course of a long simulation, one may expect *s* to fluctuate between 0 and 1 as depicted in fig. 1. Note that in multi-state systems, each metastable state A can be used as a reference to define ‘A’ and ‘not A’ states. Consequently, the two-state model translates directly to multi-state systems (appendix A).

**Fig. 1:**
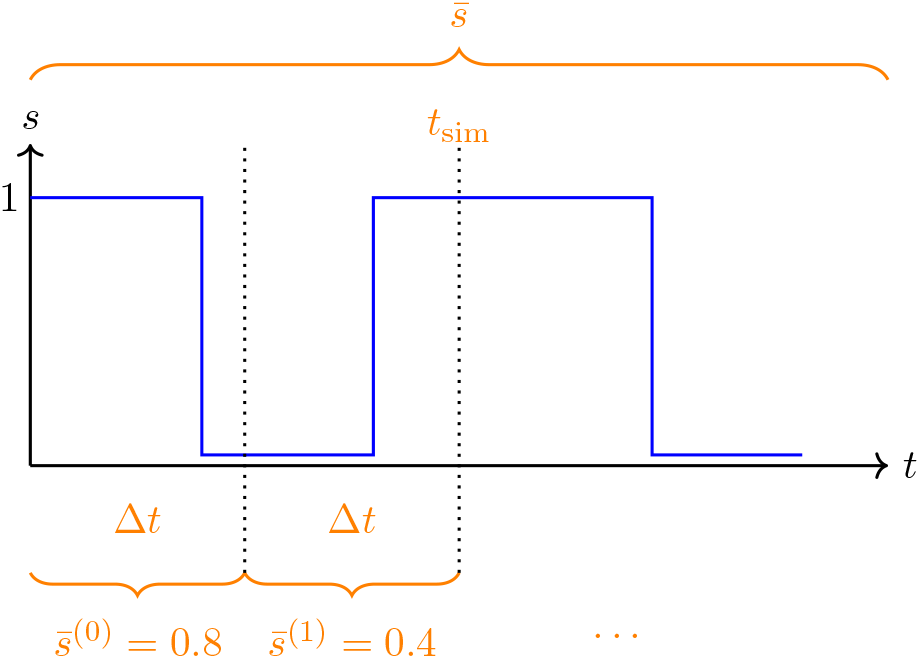
Graphical representation of the state function *s* and block averaging of a trajectory. The state variable fluctuates between 0 and 1 over the course of the simulation at any particular temperature. Δ*t* is the block size, 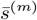 the block means, 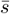 the full simulation mean and *t*_sim_ is the simulation time. The deviation of the probability estimates from each block with respect to the full simulation is 𝒮^(*m*)^.

**Fig. 2:**
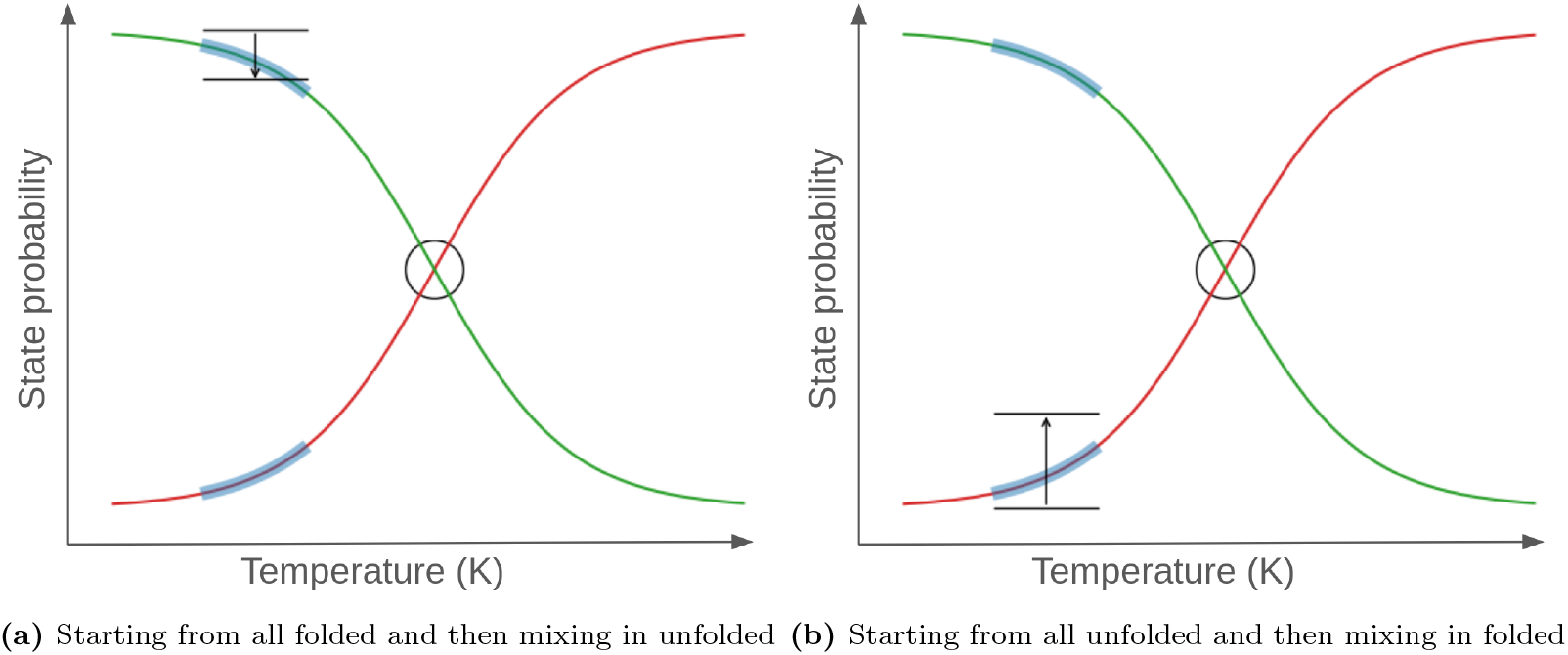
A diagram showing the shift in the probability estimates with respect to the true probability curves when the set of starting structures are changed.

To extract uncertainties from MD simulations, blocked analyses are routinely performed to address correlations in the data. Consider a simulation of length *t*_sim_, expressed in the number of time steps. Such a simulation can be divided into blocks of size Δ*t* (once more expressed in the number of time steps) such that one obtains 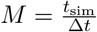 blocks, divided at points 0, *t*_1_, … *t*_*m*_ …, *t*_*M*_ in the simulation. Averages over these can be represented as

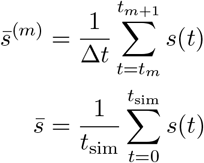

and the corresponding deviation of block means would be

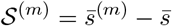

The main assumption made here is that this block-averaged deviation behaves like an OU process (see appendix A). The exponential decay of OU processes and characteristic time scales associated with them have been discussed in literature previously, see Neureither and Hartmann [21]. Hence, the assumption that such a treatment would yield results regarding the effects of the initial conditions, convergence rates and long-time behavior are not necessarily novel. Considering the difference equation for such an OU process, one arrives at the following analytical expression for 𝒮^(*m*)^:

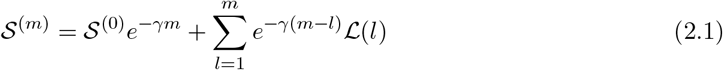

where *γ* is a friction coefficient and it is assumed that the noise term ℒ is Gaussian distributed with a zero mean and finite variance given by

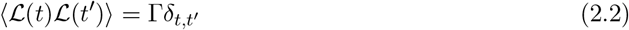

The magnitude of the noise term is determined by the size of blocks. If one has small blocks, the blocked estimates of probability of state F would vary rapidly and fluctuate between values close to 0 or 1. On the other hand, if the block size is several times the longest mean first passage time in the system, the blocked probability estimates would fluctuate very little.

### 2.2 Calculation of Standard Errors for Single Replica

The constant value 𝒮^(0)^ is the deterministic initial condition applied to the OU process. It is effectively the set of starting structures that the TREMD simulation is initialized with. The OU model provides an analytical expression for the deviation of blocked means with respect to the mean of the full simulation. To estimate the standard error in the estimator of state probabilities i.e. 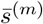, the jackknife or bootstrap methods can be applied to blocked means. One can reformulate the variance obtained from these simulations in terms of the deviation of blocked means. Combining the OU model with these reformulated expressions gives estimates of the standard error or the uncertainty in probability estimates. For more detailed derivations, appendix A.2 and section S1.2 can be referenced. However, the general form of the uncertainty when using these methods comes out to be

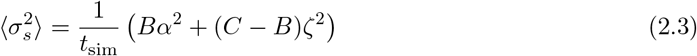

where *α* is dependent on the starting structures, i.e. *α* ≡ 𝒮^(0)^, and *ζ* is proportional to the variance Γ of the noise ℒ. The coefficients *B* ∼ 𝒪(*M*^−1^) (from where we have 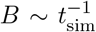 since *M* ∝ *t*_sim_) and *C* ∼ 𝒪(1) suggest that in the standard error of the mean, the effect of the choice of starting structures decays as 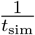, whereas the overall standard error decays as 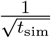. For reasonably long simulations, the effect of the initial conditions falls off faster than the overall standard error estimate.

### 2.3 Extension to Multiple Replicas

The expression above deals with a trajectory at a single temperature. This can arise either from cMD simulations or from de-multiplexed TREMD trajectories (see section 3.2). The expression for a single temperature can be reformulated as follows for ease of handling. Assuming the temperature is *T*_*i*_; *i* = 1, … *n*_reps_ and representing all temperature-dependent factors with subscript *i*,

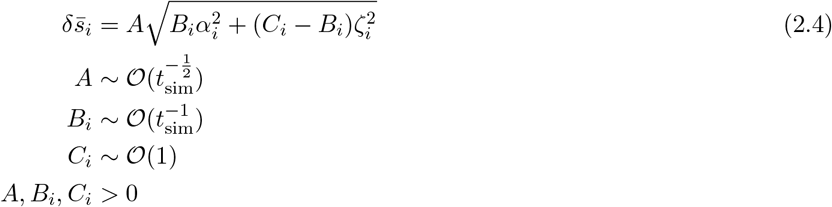

Here, we reiterate that 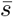 is the folded state probability. To extend the uncertainty in the estimates of state probabilities to the uncertainty in the *T*_*M*_, it is assumed that

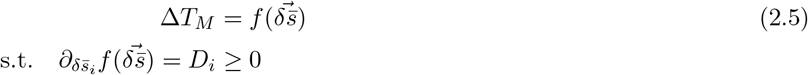

where Δ*T*_*M*_ is the uncertainty in the melting temperature estimate and *f* (·) is a non-decreasing function of the uncertainties 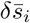 at all temperatures in the ladder. Once more, the model is tailored for a two-state system. For a multi-state system, there would be additional probability terms in Δ*T*_*M*_ for the other metastable states (each modeled as an OU process).

To determine how the uncertainty in the *T*_*M*_ estimate for a certain simulation time varies with the set of starting structures, the following procedure is adopted:

1. Consider *n*_reps_ temperatures in a temperature ladder.
2. Assume a specific set of starting structures. Let this point be represented by 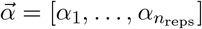.
3. Assume now that the starting structures are modified to 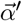. This leads to a shift 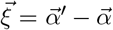.
4. Perform a directional derivative of the *f* (·) in the direction 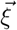 at point 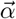.

Based on the directional derivative, it should be possible to gauge any improvement or deterioration of the melting temperature for a specific set of starting structures. Following the above process, we have

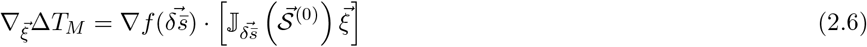

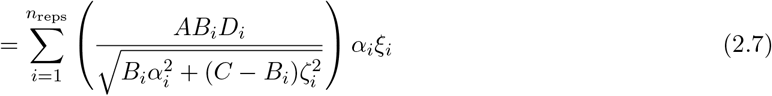

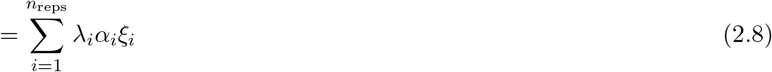

with

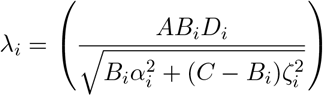

being a positive factor.

### 2.4 Observations

Consider first, temperatures below the melting temperature of the system of interest. If all replicas are initially in the unfolded state, the deviation of probability estimates from the true probability at this temperature is most likely negative within the first block. This means *α*_*i*_ *<* 0 for all *i*. Next, assume that one-third of the replicas are initialized with the folded state. In this case, the deviations may be expected to less negative, i.e. the probability estimates from within the first block would be higher. In essence, 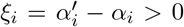. Consequently, one finds that 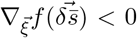. From this, it follows that for the same simulation time, the probability estimate is more accurate if replicas are initialized with a mixture of folded and unfolded starting states (as opposed to only unfolded states).

Similarly, if one initializes all replicas with the folded state, the probabilities are likely overestimated due to slow folding-unfolding dynamics. This implies *α*_*i*_ *>* 0 for all *i*. Now, if a small portion, maybe one-sixth of the replicas, are initialized with the unfolded state, the probabilities within the first block would drop. In this case, 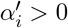 for most *i* but potentially 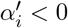 for the lowest temperatures. One can then say that *ξ*_*i*_ *<* 0 for some *i* and *ξ*_*i*_ *>* 0 for others depending on how many replicas are initialized with the unfolded state. This trend holds true for higher temperatures as well. Improvements in convergence are observed only so long as the probabilities are close to the mean expected probability over the temperature ladder. It is also worth noting that *λ*_*i*_ −→ 0 when *α*_*i*_ −→ 0. Hence, if the deviations are slightly higher or lower, the overall improvement in performance when moving either way is negligible. Effectively, the number of replicas can only be optimized to an extent; if there is enough of a mix of replicas in the initial setup, changing the starting configuration of one replica has little effect on the performance.

Based on the above arguments, it can be expected that initializing a TREMD simulation with a mixture of states is almost always more effective than using a single initial state, except if the temperature ladder is very far from *T*_*M*_.

## 3 Materials and Methods

The numerical validation of the analytical model was performed in two ways. To provide a simpler, more intuitive proof of the predictions, Parallel Tempering Markov Chain Monte Carlo (PT-MCMC) simulations were performed in an engineered double-well potential. In addition to this, all-atom explicit-solvent Molecular Dynamics simulations were performed on the fast folding protein Chignolin with three force fields. The systems, details pertaining to their simulations and subsequent processing have been discussed in this section.

### 3.1 Model Systems

The objective in both simulations is to sample the free energy landscape and make concrete statements regarding the quality and convergence of sampling. In MD, the biomolecule (Chignolin in this case) was propagated directly in phase space. On the other hand, in MCMC simulations, the free energy landscape was modeled using a double-well potential. Then, the Metropolis-Hastings algorithm was used to sample the resulting distribution.

#### 3.1.1 Model Double-Well

The constructed energy landscape, shown in fig. S1, was modeled to emulate the projection of the free energy of a two-state system onto some collective variable (such as the RMSD). For Chignolin, the projection of the free energy landscape onto the RMSD with respect to two metastable folded states is shown in fig. S3. While the fast-folding protein is actually a three-state system, it is seen that the folded and misfolded state minima merge around 400 K and the resulting profile matches the Model Double-Well (MDW) quite well.

The MDW was constructed using a linear combination of five Gaussian functions, the details of which are provided in section S2.1. In total, 8 sets of MCMC simulations were performed with 6 relatively high *β* values, i.e. low temperatures corresponding to 1 set of conventional MCMC and 7 sets of PT-MCMC. Simulations for each set of starting points were run for a total of 1.25 billion steps (250 million steps *×* 5 repetitions) per replica. This gave 7.5 billion steps per setup (1.25 billion steps *×* 6 replicas) and 60 billion steps in total (7.5 billion steps *×* 8 setups). The details of the sets of input parameters used and *β* values can also be found in section S2.1.

#### 3.1.2 Chignolin

Chignolin [22] is a fast folding protein with the sequence GYDPETGTWG. It folds into two metastable *β*-hairpin structures in water and has been studied extensively in literature. This system was chosen due to its fast kinetics, with folding times of the order of 1 *µ*s at room temperature [16, 23]. The folding-unfolding process of Chignolin as well as its Free Energy Landscape (FEL) have been studied in prior works as well [24].

The NMR structure of Chignolin with the ID: *1UAO* [22] from the RCSB Protein Data Bank [25] was used to run MD simulations. The system was prepared by solvating in an octahedral box with a molecule-to-wall distance of 20.0 Å for FF99SB [26] and 30.0 Å for FF14SB [27] and FF19SB [28]. Two Na^+^ ions were used as counter ions to neutralize the system. Subsequently, Na^+^ and Cl^−^ ions (22 for FF99SB, 45 for FF14SB and FF19SB) were added to neutralize and subsequently bring the total salt concentration to 0.2 M. Following this, energy minimization, restrained heating, slow NPT relaxation of restraints and finally NVT production runs were performed. A real space cut-off of 11.0 Å was used for the non-bonded interactions. The FF99SB force field was used for method development owing to the availability of reference values in literature. The FF19SB and FF14SB force fields were used to validate the scheme and gain further insights. It was expected that FF14SB would reproduce the experimental *T*_*M*_ more closely based on the results of simulations at 300 K performed using these force fields [11].

For the FF99SB simulations, the full temperature range of interest from 320 K to 450 K was populated with 5 temperature ladders each containing six replicas (fig. 3). For each temperature ladder, 12 sets of starting structures were tested which included different combinations of the folded, misfolded and unfolded states that Chignolin adopts with FF99SB. The folded and misfolded states have been provided in fig. S2a and fig. S2b respectively. TREMD runs of each of the 60 setups (12 starting structures and 5 ladders) were 0.95-1.0 *µ*s long. Such microsecond scale simulations were repeated 5 times for each setup, giving a total accumulated simulation time of approximately 1.8 ms (12 starting structures *×* 5 ladders *×* 6 replicas *×* 5 repetitions). It should be noted that cMD accounted for an additional 200 *µ*s of simulation time. Generally, for the results shown below, quantities (such as the standard error estimate, mean values and root mean squared errors) were averaged over the five repetitions of each setup to emulate (sub)ensemble averaging.

**Fig. 3:**
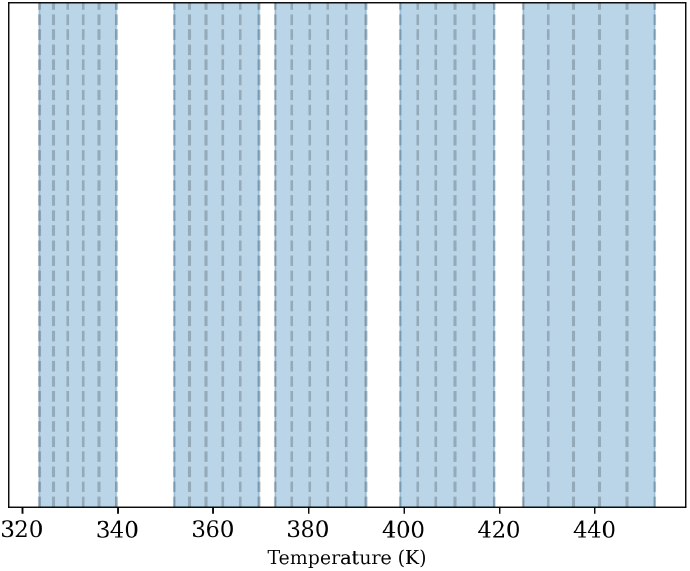
The different temperature ladders are shown as blue bands. The vertical dotted lines are the temperatures at which replicas were placed. The spacing was optimized to get an exchange success rate between 20 and 30 %. There are indexed 1 to 5 from left to right.

The simulation setups used for TREMD simulations of Chignolin are provided in table 1. For short notation, we defined a naming scheme for the TREMD setups, depending on the set of starting structures. The setups are encoded according to the pattern {*f*}f_ {*m*}m_{*u*}u, where *f, m* and *u* indicate the number of replicas in the folded, misfolded and unfolded states respectively. For example, 1f_2m_3u corresponds to 1 folded, 2 misfolded and 3 unfolded structures at the beginning of the simulation (random example). This nomenclature is used throughout this paper. Additionally, for a two-state model where only the unfolded and ‘not unfolded’ states are considered, the naming convention will be adapted to {*f*}f_{*u*}u where ‘f’ stands for the ‘not unfolded’ state.

**Table 1.** Shows the names of 12 setups and the number of replicas in the ladder starting from each metastable configuration. Total times were calculated for 5 ladders *×* 5 repetitions (per ladder). This means simulation time per ladder, per repetition of starting structure set is 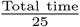.

| Setup | # F | # M | # U | Total time (μs) |
| --- | --- | --- | --- | --- |
| 1f_1m_4u | 1 | 1 | 4 | 25.0 |
| 2f_3m_1u | 2 | 3 | 1 | 25.0 |
| 0f_0m_6u | 0 | 0 | 6 | 24.9 |
| 0f_2m_4u | 0 | 2 | 4 | 24.9 |
| 0f_4m_2u | 0 | 4 | 2 | 24.8 |
| 0f_6m_0u | 0 | 6 | 0 | 24.9 |
| 2f_0m_4u | 2 | 0 | 4 | 24.8 |
| 2f_2m_2u | 2 | 2 | 2 | 24.9 |
| 2f_4m_0u | 2 | 4 | 0 | 24.9 |
| 4f_0m_2u | 4 | 0 | 2 | 24.9 |
| 4f_2m_0u | 4 | 2 | 0 | 25.0 |
| 6f_0m_0u | 6 | 0 | 0 | 24.9 |

### 3.2 Melting Temperature Extraction

Constant temperature trajectories were obtained either using conventional MD simulations at multiple temperatures or by de-multiplexing replica exchange simulations (TREMD or PT-MCMC). For the latter class of simulations, de-multiplexing generated thermodynamically continuous trajectories from structurally continuous trajectories.

#### 3.2.1 Representative Structures of Metastable States

For Chignolin, reference structures were identified via Agglomerative clustering using average linkage, a cut-off of 2.8 Å and the 2D-RMSD of heavy atoms as the precomputed similarity matrix. We followed an approach similar to the one described by Shao et al. [29]. The identified structures are shown in fig. S2. The heavy-atom RMSD of the MD trajectories with respect to the reference structures were then computed. For each RMSD, histograms were constructed and the first minimum was calculated from a contour estimated with Gaussian Mixture Models. The minima were used as cut-offs for the corresponding reference structures.

The metastable state with respect to which the RMSD was minimum and below the associated cut-off, was assigned to every frame. If a frame had RMS distances greater than all stable state cut-offs, then the unfolded state was assigned to the frame (even when the configuration was persistent for non-negligible windows of time). For the MCMC simulations, the process was much simpler: the barrier peak was identified and all trajectory points to the left of the barrier were assigned to the folded state and those to the right were assigned to the unfolded state.

#### 3.2.2 State Probability and Melting Temperature Estimation

With the state of each frame identified, state arrays were constructed using one-hot encoding. The TREMD simulations were analyzed using a two-state (one state array for ‘not unfolded’ i.e. folded or misfolded) and a three-state (separate state arrays for folded and misfolded states) model, giving three melting temperatures. Averaging these state arrays yielded temperature-dependent probability estimates for each state. The resulting probability curves were then processed using the van’t Hoff analysis and melting temperaturess were extracted for each metastable state. We followed an approach similar to the one described by Quoika et al. [30] for a temperature-dependent conformation change of thermosensitive polymers.

### 3.3 Uncertainty Estimation

The analyses were designed with two primary objectives: (i) to examine how the choice of starting structures for replicas influences the convergence of small-ladder TREMD, and (ii) to assess the utility of small-ladder TREMD simulations for estimating the *T*_*M*_ of small biomolecules. Accordingly, the first set of analyses focused on state-probability estimates and were used to evaluate the mathematical model developed in section 2 and section S1. Hence, they were used to assess the impact of initial conditions on the precision and accuracy of probability estimates even in fast-folding systems such as Chignolin. The second set of analyses focused on *T*_*M*_ estimates as a direct measure of small-ladder TREMD simulation performance.

#### 3.3.1 Trajectory Blocking and Probability Estimation

Consider first, a single setup defined as a combination of temperature ladder and starting structures (e.g., ladder 1 initialized with 2f_2m_2u). For each of the six replicas, trajectories were first blocked. For the PT-MCMC simulations, it was assumed that 10,000 frames (1 million MCMC steps) were sufficiently long to decorrelate the data. The process was slightly more involved for TREMD simulations.

To determine an appropriate block size, the RMSD time series at the lowest temperature of each ladder was first analyzed to estimate the autocorrelation time. This autocorrelation time was then used as the lag time for constructing a Markov state model (MSM) with PyEMMA, performed separately for each repetition of each simulation setup. From each MSM, mean first passage times (MFPTs) were computed, and the largest MFPT was taken as a rough estimate of the minimum block size required for that repetition of the TREMD simulation setup. For each setup, the largest MFPTs (defined above) were averaged over the five repetitions. The minimum of these averaged values for each ladder was selected as the block size to ensure a sufficient number of statistically independent blocks. Finally, these values were clipped to ensure a minimum block size, giving 20 blocks for ladder 1 simulations and 38 for the other ladders (since the calculated block sizes for those setups fell below the cut-off of 10 ns per block). The minimum was set since extremely low block sizes resulted in artifacts during post processing.

To derive statistics for the probability estimates, resampling was performed in three complementary ways. First, bootstrapping from the first *m* blocks; second, subsampling with replacement from all *M* blocks; and third, subsampling with replacement excluding the equilibration phase i.e. from the latter *M* − *k* blocks, where *k* is the number of equilibration blocks.

These different resampling schemes are explained in more detail in section S2.3. Once resampling was performed, the mean and variance were calculated for the constructed datasets to determine statistics for the state probabilities. To illustrate the effect of the equilibration phase, we investigated the mean standard error estimates of probabilities with and without this initial period. Depending on the starting conditions, the impact varied considerably. Since fast and slow time scales could not be effectively separated during the error analyses, quantitative insights regarding how far the equilibration phase extends could not be made. However, strong tendencies could be observed based on the shift of the constants of proportionality between the same setups with and without the initial blocks.

#### 3.3.2 Reference Values for State Probabilities and Melting Temperatures

To compare different simulation setups, reliable reference values were necessary. Since we had accumulated large amounts of data, we extracted references for the FF99SB force field from our simulations. It was seen that removing the first 250-300 ns of simulation time generally gave equilibrated results at the lowest temperatures (see fig. 6 for details). The equilibration phases were removed from all TREMD simulations and the corresponding probabilities per temperature were calculated as an average over 12 setups and 5 repetitions for 30 temperature points (5 ladders and 6 replicas each). This gave a total of 42 *µ*s per temperature or 1.26 ms of simulation time overall. The van’t Hoff analysis of these simulations is provided in fig. 4.

**Fig. 4:**
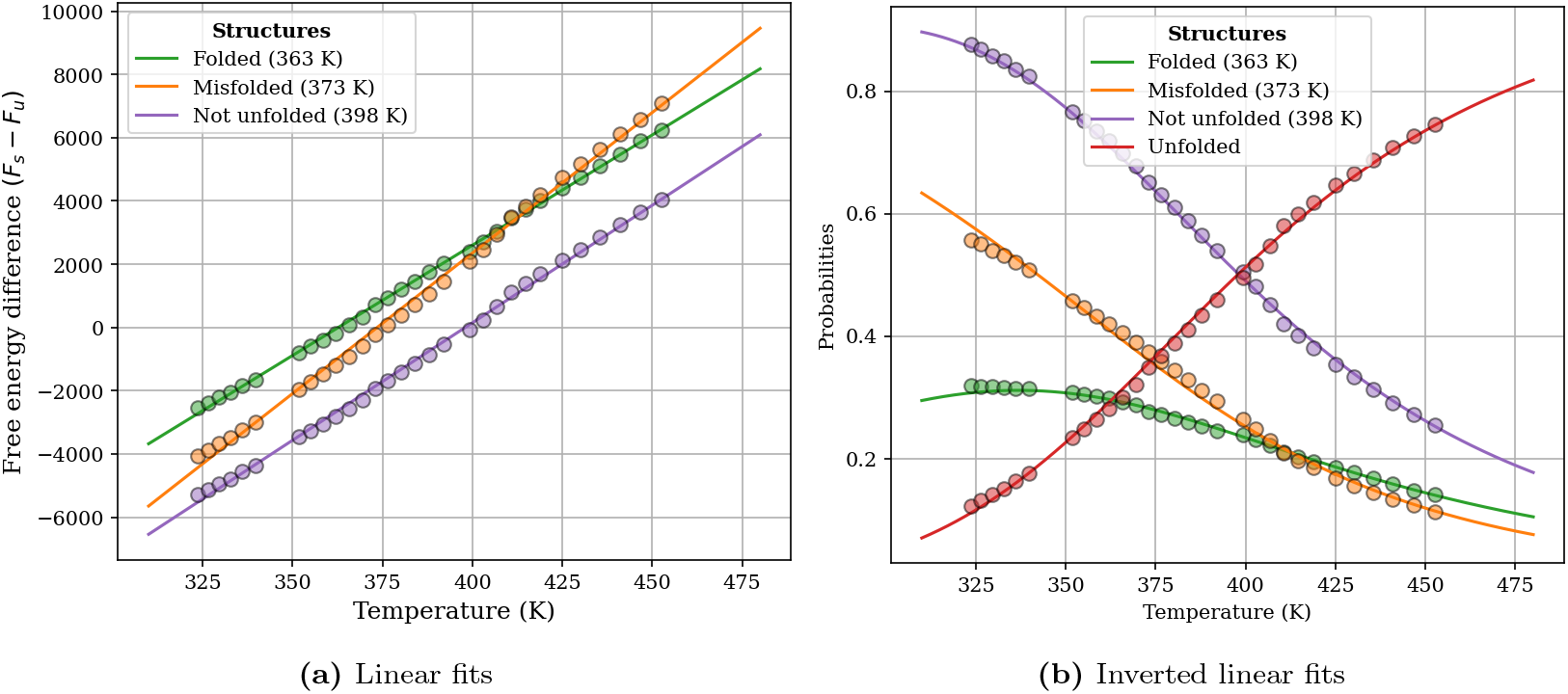
The probability estimates per temperature was obtained by averaging over 12 setups and 5 repetitions of TREMD simulations excluding the first 250-300 ns per setup, per repetition. Similar analyses for cMD have been provided in fig. S4. Once the free energy differences ΔF were calculated with the van’t Hoff analysis, melting curves were generated using the reverse process i.e. *p* = −*k*_*B*_*T* ln *F*.

Hence, reference distributions were constructed by pooling approximately 1.26 ms of equilibrated TREMD data. Following the variance of state-probability estimates, the accuracy (root mean-squared error or RMSE) of mean bootstrapped state-probabilities with respect to reference probabilities was also computed. Combined, the variance and accuracy helped distinguish between imprecise and inaccurate, precise and inaccurate, imprecise and accurate and precise and accurate probability estimates.

#### 3.3.3 Resampling for Melting Temperature Estimation

Since the ‘^*m*^*C*_*m*_’ (this is intended for notation purposes – in standard combinatorics it evaluates to 1) or bootstrapping strategy is most likely to be used by practitioners, melting temperatures were obtained using the bootstrapped probability estimates. Box-plots were prepared for these bootstrapped *T*_*M*_ estimates and provided summaries of the setup-dependence of the performance of small-ladder TREMD simulations. In this case, the median and inter-quartile range (IQR) were used to describe final estimate and uncertainty because of large deviations from measures of central tendency. Unlike probabilities, which are constrained to [0, 1], log-probabilities or free-energy estimates are unrestrained. Hence, outliers tend to be far from the median or mean, in extreme cases extending to unphysical regimes.

## 4 Results and Discussion

First, PT-MCMC simulations performed in the MDW are discussed using the schemes described in section S2.3.2 and section S2.3.3. Then, the MD simulations are discussed in great depth from the perspective of the probability estimates and melting temperatures, their standard error and root mean-squared error (precision and accuracy).

### 4.1 Minimal Model

The MCMC trajectories were processed as described in section 3.2 and section 3.3. The mean number of replicas in the folded state over the whole ladder can be seen for the Markov Chain Monte Carlo simulations in fig. 5.

**Fig. 5:**
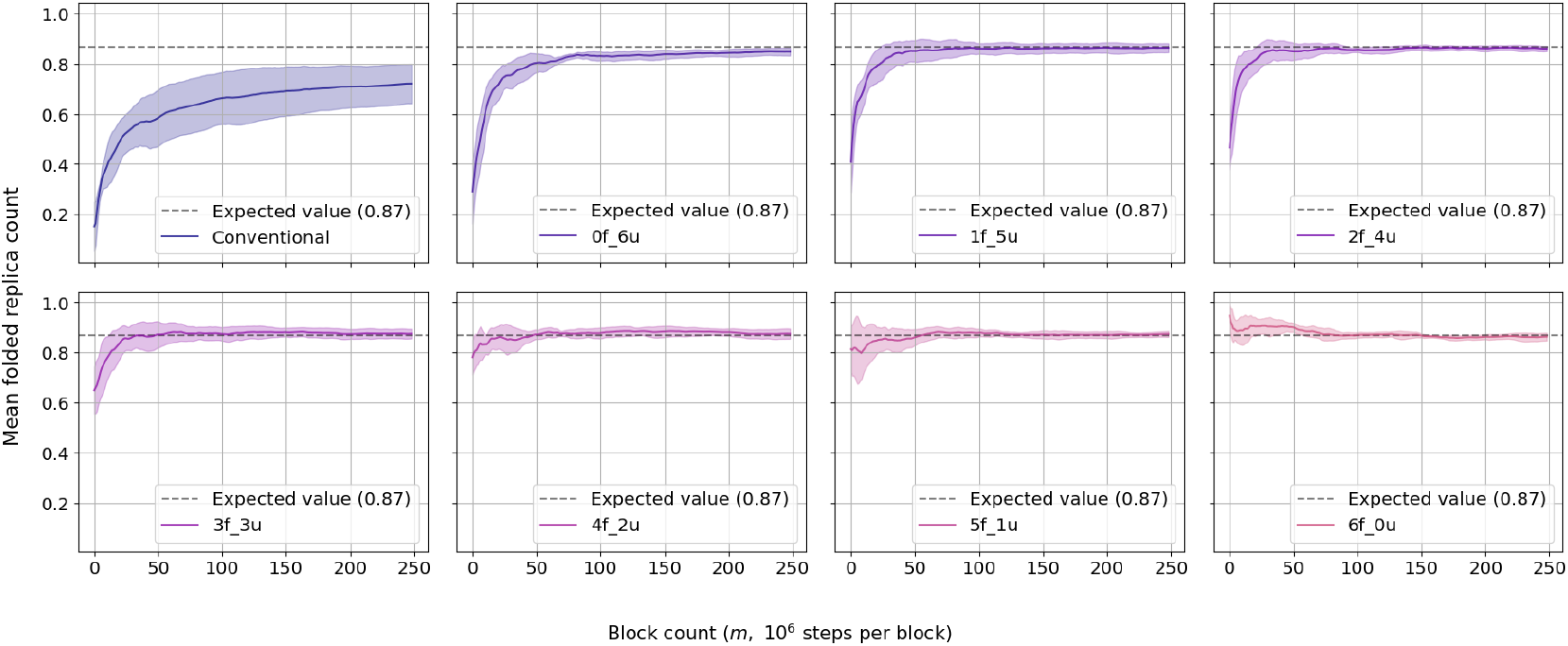
The mean number of replicas in the folded state for the bootstrapped (^*m*^C_*m*_) case of the MCMC simulations. The ideas seen in Rosta and Hummer [18] have been validated and there is a clear discrepancy between the length of the equilibration phase for different setups. The equilibration phase was identified to be approximately 35 % of the total block count *M* based on where the curves approach their expected value. These numbers were obtained by averaging the bootstrap averaged folded state probability of each replica over 5 repetitions.

From fig. 5, it is intuitively clear that the simulations equilibrate to the expected distribution of replicas per state. However, the effect of the initial conditions wanes over different scales. As expected, conventional MCMC (i.e., without replica exchange) performs the worst. The setups 3f_3u, 4f_2u, 5f_1u perform well when judging on the basis of the mean folded replica count, as predicted by the theoretical model (section 2, applies since the replica count per state is linearly dependent on state probabilities). Additionally, it was also expected that 5f_1u would perform the best. This is evident from fig. 5, since this setup begins from the highest mean count of replicas in the folded state (albeit marginally). Overall, the toy systems bolsters the claims made on the basis of the theoretical model.

To further visualize the performance, we defined the proportionality constant

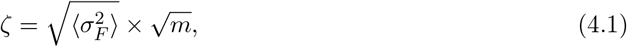

where 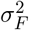 is the variance estimate of the folded-state probability *p*_*F*_ obtained by resampling *m* blocks with replacement. The values of *ζ* for all three sampling schemes described in section S2.3 are shown in fig. S8. Their relative values have been reported in fig. S7. Estimates of the *T*_*M*_ of the MDW from PT-MCMC initialized with different starting structures is shown in fig. S6. It is evident from fig. S7 and fig. S8 that the constants of proportionality change the least when replicas are initialized predominantly in the folded minimum (left of the barrier). The shift in precision observed (fig. S7) when the initial conditions drift away from the optimum can be attributed to the bias introduced by the initial conditions. In the context of the OU model introduced above, this bias increases the tendency to sample points from the tail ends of the Gaussian noise distribution ℒ. When these blocks are resampled, the estimate of the overall uncertainty quantified by *ζ* increases.

The reader may note from fig. S6 that the accuracy of the *T*_*M*_ estimates obtained for the 1f_5u and 2f_4u cases (high ratio of unfolded replicas initially) are as high as that of 5f_1u beyond 100 million steps (100 blocks). This must however, be taken within the context of higher precision of the 5f_1u simulations for lower time scales. Similar observation is made for MD simulations, which form the next topic of discussion.

### 4.2 Chignolin - FF99SB

As mentioned in section 3.1.2, there were 60 possible setups, i.e. combinations of the temperature ladders and sets of starting structures. Here, we focus on only the most relevant cases – namely 6f_0m_0u, 0f_6m_0u, 0f_0m_6u, 2f_2m_2u, 2f_3m_1u and_1f_1m_4u simulated with the lowest two temperature ladders. These correspond to approx. 325-340 K (ladder 1) and 350-370 K (ladder 2), as shown in fig. 3. The first three starting structure combinations are edge-cases (all replicas in one state) and the next three contain a mix of all three metastable states initially. Occasionally, other setups will be referenced to provide more concrete proof of claims.

#### 4.2.1 State Probabilities and the Effect of Initial Conditions

Rosta and Hummer [18] predict that the TREMD ladder oscillates about an equilibrium distribution of replicas in metastable states. This effect was observed in all simulations performed in the present study. Consider fig. 6, where running average of the number of replicas in each of the three metastable configurations (averaged over five repetitions) has been shown for ladder 1. An equilibrium distribution is observed in nearly all cases within 1 *µ*s, regardless of the initial conditions. However, in several cases the equilibration phase is pronounced, indicating that initial conditions are often non-negligible at low temperatures.

**Fig. 6:**
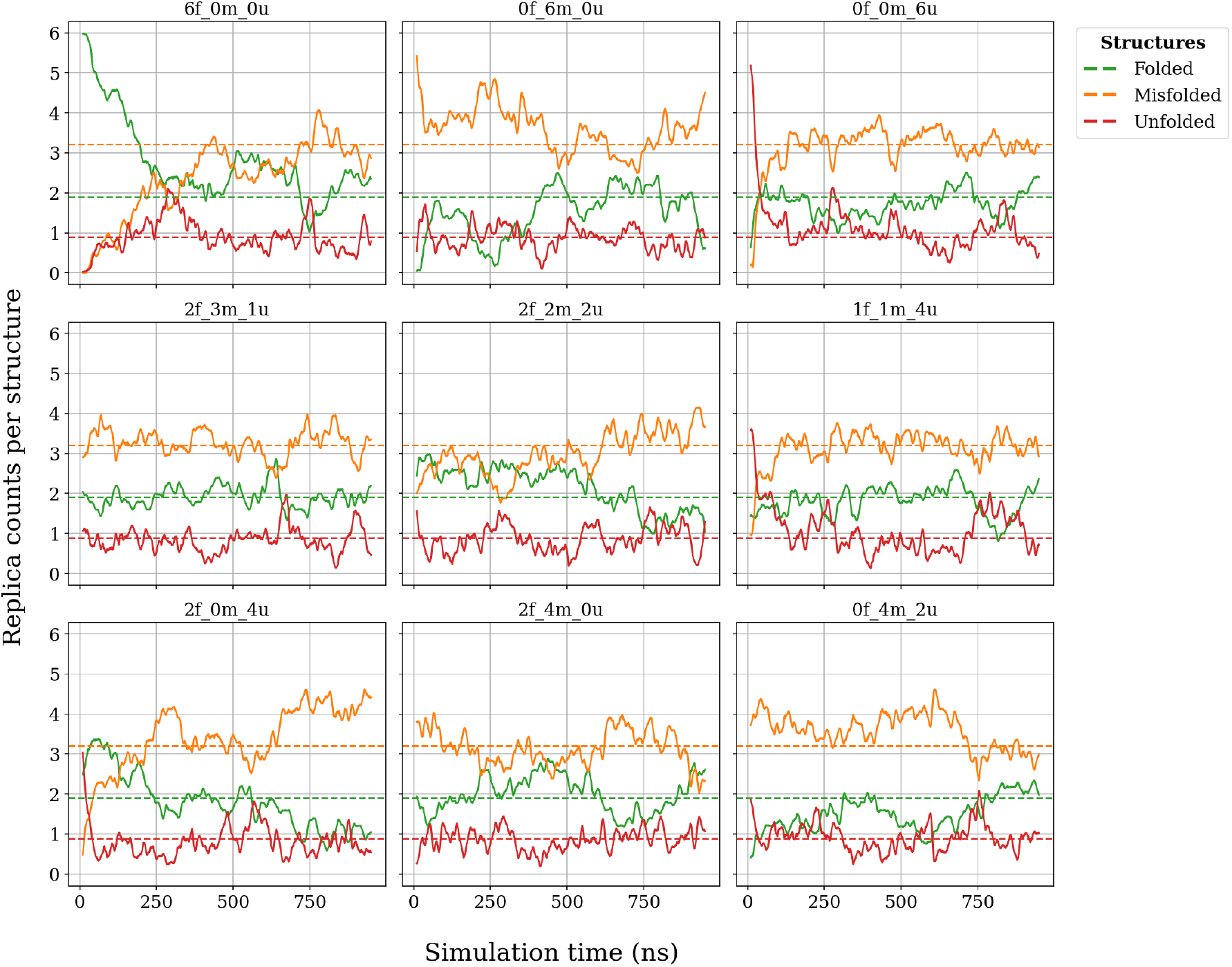
Running average over 100 frames of replica count per structure for temperature ladder 1. For each setup, the time series of replica counts was computed for all five repetitions and then averaged. The running average was calculated from this ensemble averaged time series. Horizontal guide lines indicate the expected replica counts for FF99SB, obtained from fig. 4 (mean probability per state times replica count). In most cases shown, a distinct equilibration phase is observed. This region is vanishingly small for the 2f_3m_1u setup, as expected.

For temperature ladder 1, the expected distribution 2f_3m_1u quickly achieves the optimal replica count per metastable state. The setup 2f_4m_0u simulated with temperature ladder 1 produces fairly fast equilibration similar to 2f_3m_1u. This is in agreement with the conclusions of the OU model in section 2.4 and with the PT-MCMC simulations in section 4.1.

Similar plots for all setups can be found in fig. S14 and fig. S15 and it can be seen that simulations at higher temperatures equilibrate faster. Since the folding kinetics of Chignolin follow Arrhenius behavior when a two-state assumption is made (as done in van der Spoel and Seibert [16]), there is some justification for this observation. It can therefore be argued that, starting structure optimization does not significantly improve convergence behavior at high temperatures. For systems with non-Arrhenius kinetics (such as nucleic acid hairpins, Wallace et al. [31]), higher temperatures may also benefit from the optimization of starting structures.

Additional proof of the improvement in performance when initializing TREMD with the optimal distribution of starting structures can be seen in the standard error of state probabilities estimated with the bootstrap scheme, fig. 7. We also performed the same analysis with the jackknife scheme and found completely analogous results. The jackknife and bootstrap errors for all setups are shown in figs. S16 to S20 and figs. S21 to S25. Comparing different starting structures and temperature ladders, it is visible that the uncertainty (standard error estimate) in the probability estimates is least for the expected distribution of replicas (per metastable state) for the lowest temperature ranges.

**Fig. 7:**
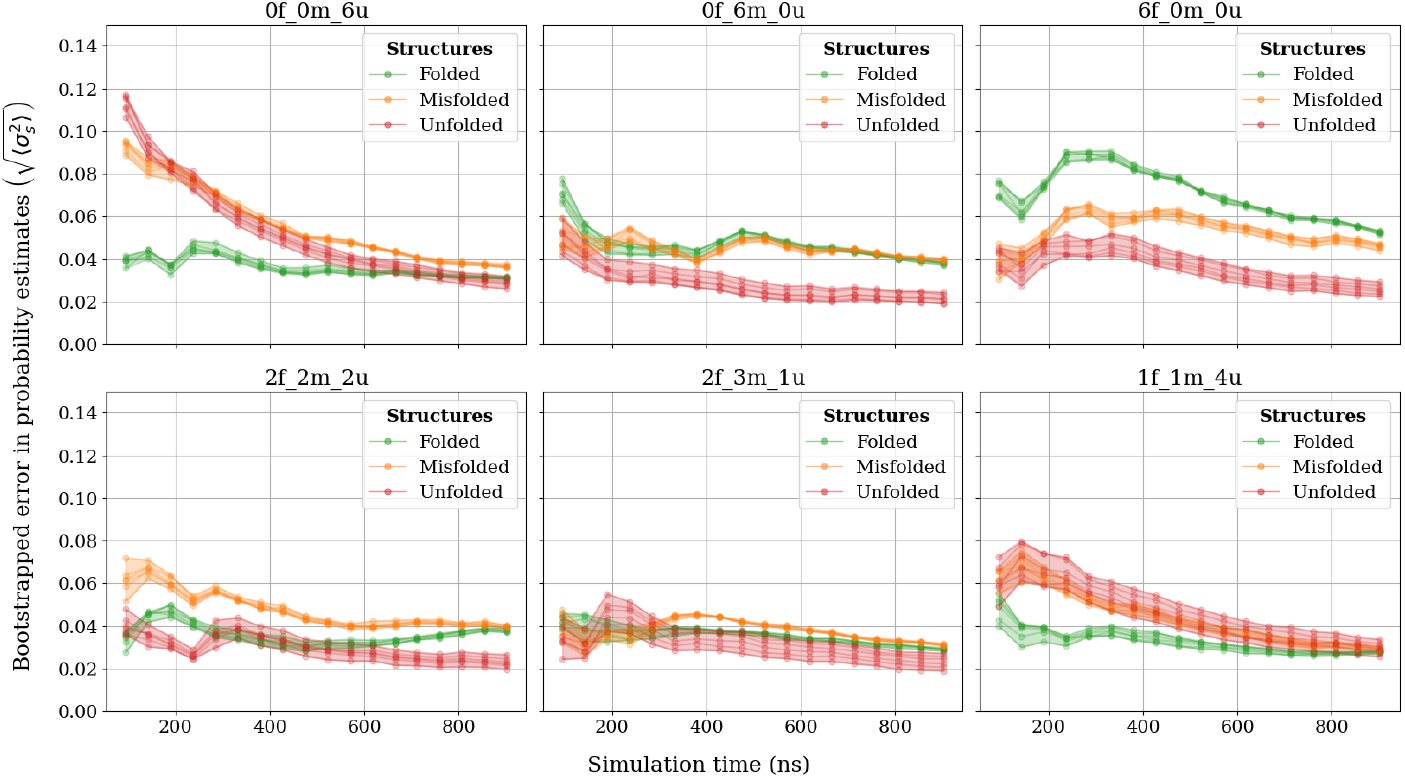
Bootstrap error of ladder 1 simulations averaged over five repetitions of each setup. All six replicas have been included. It is clear that lowest average uncertainty in probability estimates is achieved for the optimal setup 2f 3m 1u. However, this improvement is marginal and setups such as 4f_2m_0u and 0f_2m_4u also have low uncertainties for the estimated probabilities.

Uncertainty estimates based on the bootstrap variance are insufficient for identifying the optimal setup, as they do not distinguish between precise but inaccurate and precise and accurate results. Therefore, accuracy is considered next and is shown in fig. 8 for temperature ladder 1. It is evident that the 2f_3m_1u setup does not yield more accurate results than other setups; rather, its accuracy is arguably lower than 0f_0m_6u after approximately 1 *µ*s. This trend is also observed for other setups in figs. S26 to S30 for the corresponding optimal distributions (of replicas in metastable states). The utility of optimizing initial conditions is therefore greater for short simulations. For the expected replica count per metastable state, the accuracy is nearly constant for the full duration of the simulation. This should not come as a surprise as the *T*_*M*_ estimates themselves depicted in figs. S9 to S13 are stable for much of the simulation time.

**Fig. 8:**
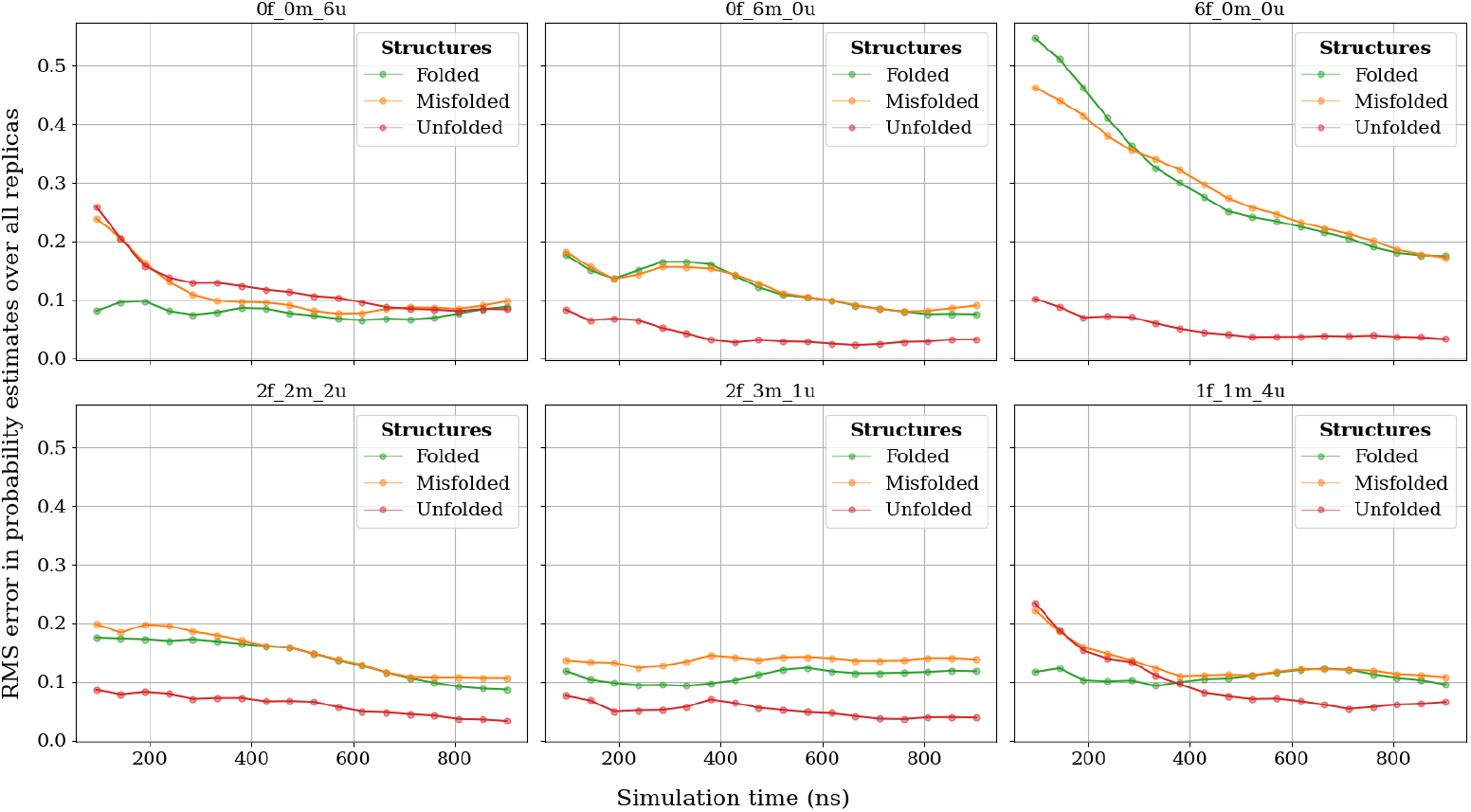
RMSE of bootstrapped probability estimates averaged over five repetitions for ladder 1 simulations. It can be seen that the worst-case setup (6f_0m_0u) is more than 5 times slower than the relatively well-initialized setups (2f_3m_1u, 2f_4m_0u).

From the high temperature simulations, it followed that adjusting the choice of starting structures becomes less important as the kinetics become faster. However, increasing temperature need not lead to faster kinetics. In general, a mixture of states should be preferable to initialization from a single state, provided all the metastable states are known. When no prior information is available, high-temperature simulations with all replicas initialized in the unfolded state provide the most favorable balance between accuracy and precision for short simulations.

Simultaneously, it must be noted that even the unimportance of initial conditions at high temperatures for systems such as Chignolin are not strong enough reasons to simulate only at those temperatures. If one considers the extrapolation from high temperatures to estimate melting temperatures shown in figs. S12 and S13, for example, it is clear that the flatness of the probability curves yields inaccurate and imprecise results. Similarly, the flipped stability of the folded and misfolded states at very high temperatures with the FF99SB force field can yield qualitatively incorrect values for melting temperatures upon extrapolation (see fig. 4).

Overall, the combination of short ladders and optimized initial conditions should quickly provide good estimates of the state probabilities for small molecules. Generally, a mixture of states performs better than single states for starting structures for short simulations, which should be evident from the bootstrapped standard error estimate and RMSE plots. Accurate and precise estimates of the probabilities need not translate directly to the same qualities for melting temperatures. Ladders shifted closer to the *T*_*M*_ provide better estimates and interpolation between 2 or more small ladders improves the reliability of results considerably. This is discussed in more detail in section 4.2.2.

#### 4.2.2 Small Ladder TREMD for Melting Temperature Estimation

Consider fig. 9a, which shows the melting temperatures obtained by extrapolating simulations from the lowest temperature ladder. The 6f_0m_0u setup produces qualitatively incorrect results for FF99SB, indicating that unfolding times from this state are potentially very high. Almost all setups (excluding those beginning with more than 50% replicas in the folded state) reproduce the correct relative stability of the metastable states of Chignolin after ∼ 1 *µ*s (see fig. S9) even when extrapolating ladder 1. After 1 *µ*s, the setup 0f 0m 6u produces melting temperatures close to the values generated by 2f 3m 1u, which is the expected optimal setup. Simultaneously, after 1 *µ*s, 1f_1m_4u also generates accurate *T*_*M*_ estimates for all states relative to the reference values from fig. 4. However, it should be noted that the least variation of *T*_*M*_ estimates between short and long simulation times is for 2f_3m_1u. This supports our claim that the optimization of initial conditions is most beneficial for short simulations.

**Fig. 9:**
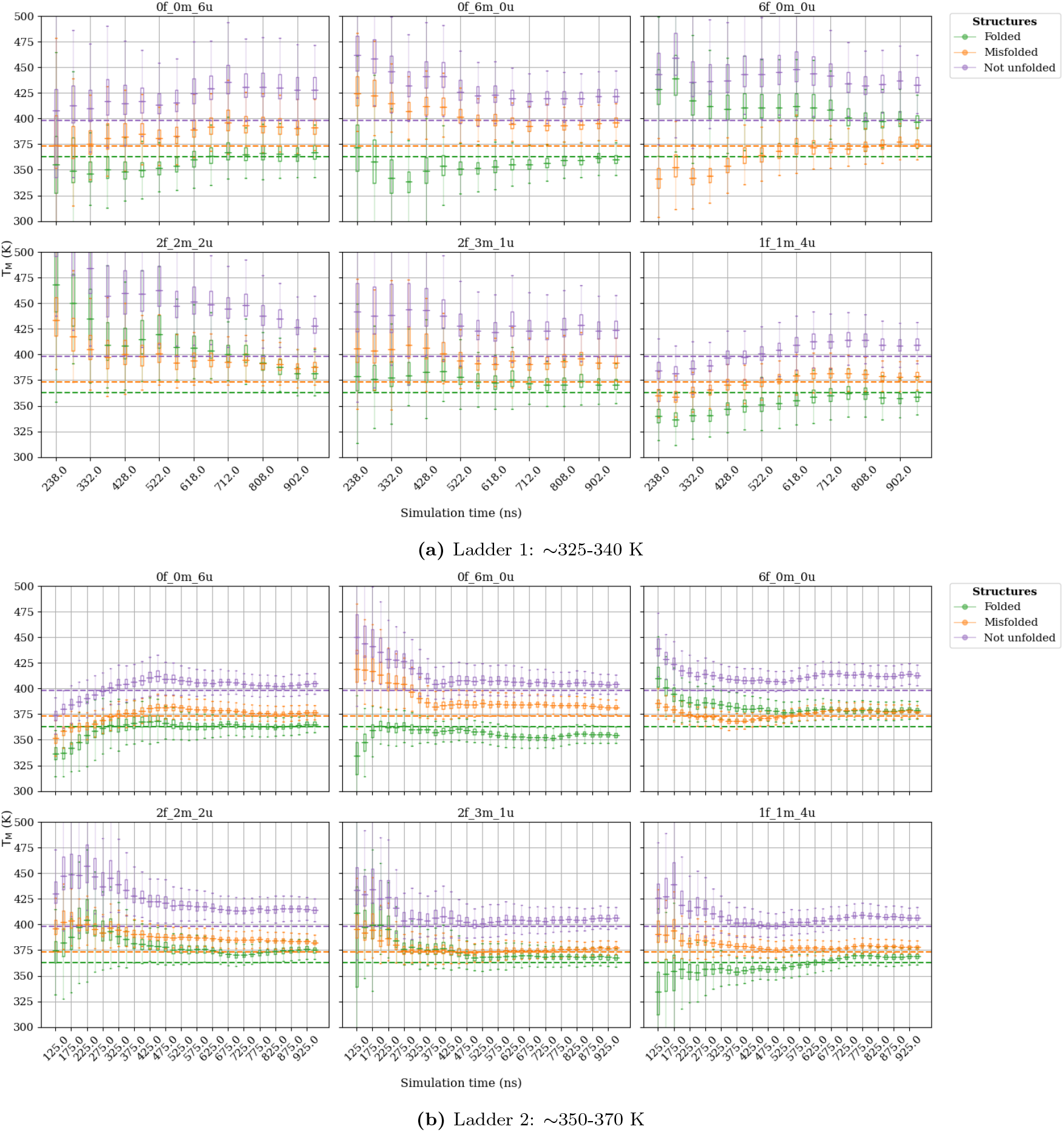
Bootstrapped *T*_*M*_ estimates from extrapolation of ladder 1 and 2 simulations. The dotted lines show the reference melting temperatures for folded, misfolded and ‘not unfolded’ states obtained from fig. 4. Ladder 1 simulations have fewer blocks due to larger effective first passage times between states (see subsection 3.3).

Temperature ladder 2, the results for which are shown in fig. 9b, produce significantly improved *T*_*M*_ estimates. It seems that initialization with a high proportion of the unfolded state typically tends to return more accurate results for simulation times ∼ 1 *µs*, as is visible from 0f_0m_6u, 1f_1m_4u, 2f_0m_4u and 0f_2m_4u (see fig. S10 for all setups). Here as well, the 6f 0m 0u setup produces qualitatively incorrect results even after 1 *µ*s. Similarly, 0f 6m 0u overestimates the misfolded state probability as expected. The inaccuracies in 2f_3m_1u and 2f_2m_2u with ladder 2 for short simulation times potentially arose as a result of folding event(s) combined with asymmetric transition times (slower unfolding than folding). When considered in the context of preset starting structures, such folding would lead to a fast deviation from the expected distribution of replicas. However, such rare instances of folding-unfolding may also occur when a high proportion of unfolded replicas is used and so is not a reason to discount starting structure optimization.

From figs. S9 to S13, it is apparent that the further away the temperature ladder is from the melting temperatures of metastable states, the less accurate the estimates obtained by extrapolating from a single ladder become. From ladder 4 simulations (fig. S12), for example, one sees that extrapolation gives folded state *T*_*M*_ estimates that are imprecise in almost all cases. Extrapolation from ladder 5 simulations (fig. S13) consistently underestimates all melting temperatures. Single ladders only yield accurate estimates of *T*_*M*_ if they cover the exact temperature window of interest (fig. S11). This would be highly difficult to achieve for unknown systems.

High temperature simulations equilibrate faster than low temperature simulations. However, extrapolation can be unstable for *T*_*M*_ estimation. Combining data from multiple small temperature ladders via interpolation can have an ‘anchoring’ effect and prevent instabilities during the van’t Hoff analysis. Each ladder would encode the probabilities of the states in the corresponding temperature regimes. Based on these observations, interpolation could yield accurate melting curves. If no prior information about the *T*_*M*_ of the system is available, simulations should first be set up at high temperatures. What temperature range is considered to be high needs to be decided based on knowledge about the thermal stability of the protein and general performance of the force field (section 4.3). The unfolded state appears to be the most reliable starting structure at high temperatures when no prior information is available, from fig. 9.

Since there are ^60^C_2_ = 1, 770 possible combinations of setups when taken in pairs in this study, the discussion below focuses on demonstrating the effectiveness of method through a plausible example. Consider a scenario where ladder 5 simulations are initialized 0f 0m 6u. From fig. S13, if simulations are stopped before 300 ns, the resulting *T*_*M*_ estimates would lie below temperature ladder 2. After 300 ns however, the melting temperature estimates would largely overlap with temperature ladder 2. To refine the *T*_*M*_ values, subsequent simulations could then be performed with either ladder 1 or ladder 2 depending on the length of the initial simulations. The set of starting structures can also be adjusted on the basis of the probability distribution obtained from the 0f 0m 6u, ladder 5 simulations. The results of interpolation between ladder 5 and ladder 1 or 2 with different starting structures have been provided in fig. 10. Here, the ladders correct for each other, giving accurate and precise *T*_*M*_ estimates in most cases. Noticeably, 2f_2m_2u ladder 2 (which overestimated the *T*_*M*_ during extrapolation) experiences an improvement in precision when interpolating with high temperature simulations. The estimates also appear much more stable, with deviations only of the order of 5 K over the entire duration of the simulations.

**Fig. 10:**
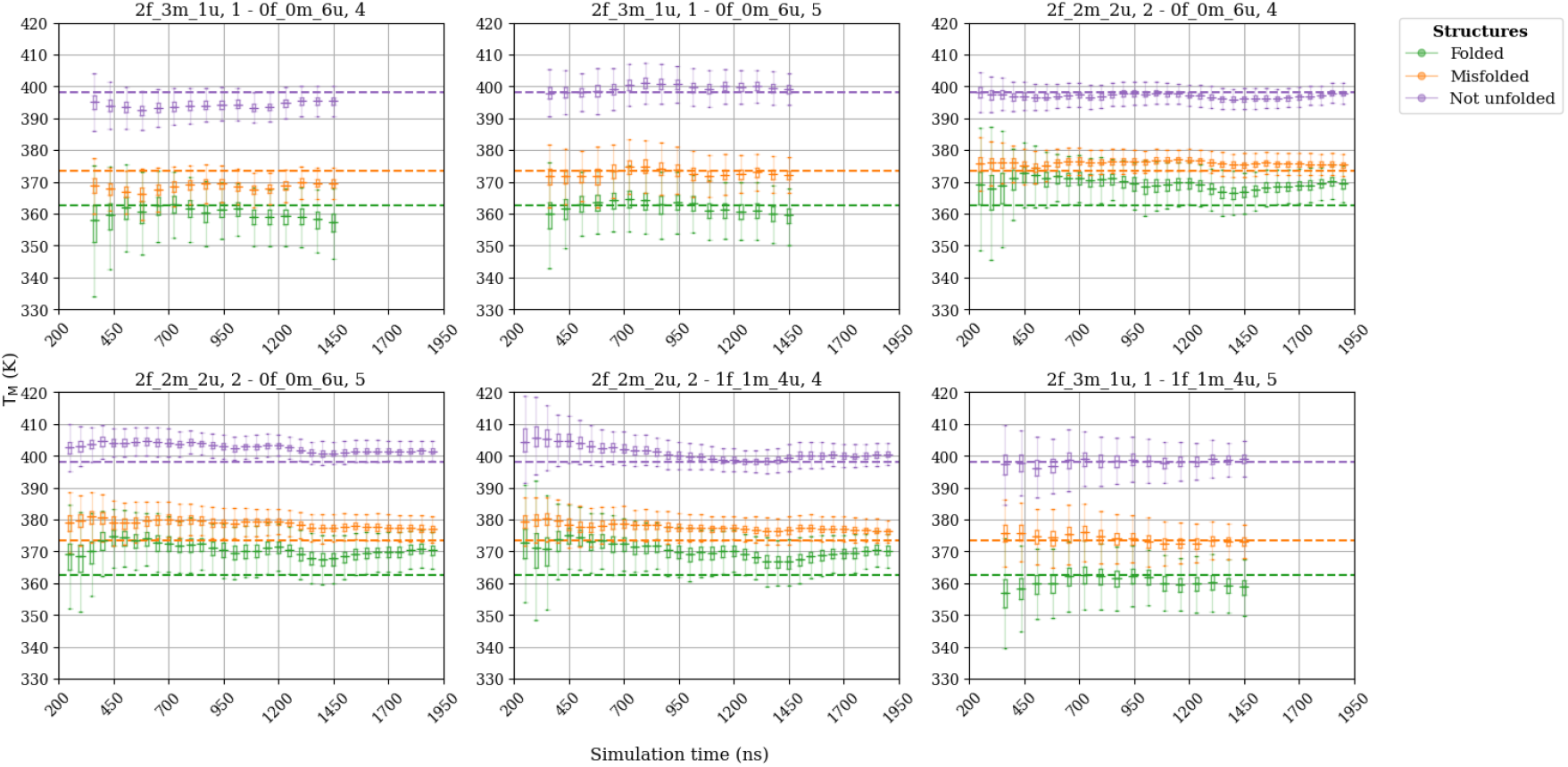
Bootstrapped *T*_*M*_ estimates from interpolation between different ladders. The dotted lines show the reference melting temperatures for the folded, misfolded and ‘not unfolded’ states obtained from fig. 4. Interpolation between small ladders with carefully chosen initial conditions provides highly accurate *T*_*M*_ estimates. The X-axis gives the sum of simulation time per block for both ladders combined. There are fewer points when interpolation is performed with ladder 1 because blocks were 47.5 ns long as opposed to 25 ns for other ladders.

An approach where small ladder TREMD simulations are first performed at high temperatures and subsequently shifted to lower temperatures with iteratively adjusted starting structures is highly viable for small molecule *T*_*M*_ calculations. These simulations would equilibrate faster and can be used when resources are limited. The full temperature range of interest need not be populated with replicas - smaller sections of the melting curve can be reproduced by starting from high temperatures and slowly sliding the ladder down. The recommendation for starting from high temperatures originates from the observation that kinetics are generally faster there. Consequently, simulations can be stopped at ∼100 ns to get the initial estimate of melting temperatures (see fig. S13 for ladder 5 simulations). It should also be clear that low temperature ladders introduce noise and make *T*_*M*_ estimates imprecise when taken alone. However, both high and low temperature regimes are necessary to generate accurate and precise melting temperatures, as they anchor each other and prevent instability during post-processing. The latter particularly benefits from the optimization of starting structures.

### 4.3 Chignolin Simulations with FF14SB and FF19SB

In this section, some results from the simulation of Chignolin with FF14SB and FF19SB using the small ladder formulation presented above have been provided. The simulations with these force fields were performed using only a single set of starting structures – 2f_2m_2u. They were performed in two temperature regimes. One close varying from 355 K to 370 K and the other varying from 323 K to 333 K. The ladders used 6 replicas spaced to achieve exchange success rates in the 20-30 % range. Analyses were performed as described in section 3.2 and section 3.3 with a block size of 9.5 ns.

The extrapolated results for the simulations with FF14SB have been provided in fig. S31. The combined (interpolated) results are shown in fig. S32. The estimates, particularly those of the (natively) folded and ‘not unfolded’ states are close to the experimental *T*_*M*_ of 310 K-315 K found by Honda et al. [22]. It is seen that taken alone, the higher temperature ladder provides highly imprecise estimates of the *T*_*M*_ for the folded and ‘not unfolded’ states. The result is imprecise and inaccurate for the misfolded state. This can be attributed to the slow variation of the probability curves at those temperatures. If one has a flat segment of the melting curve, the information encoded is too little to meaningfully extrapolate (see previous section and section 3.3.3 for further discussions). The low temperature ladder provided accurate but imprecise results. However, when combined, they provide relatively accurate and precise results.

The same problem of high temperature ladders was observed in the FF19SB simulations as well, for which the extrapolated and interpolated *T*_*M*_ estimates have been shown in fig. S33 and fig. S34 respectively. The extrapolated results from the low temperature ladder have also been provided in table 2. The high temperature ladder and interpolation generated unphysical or qualitatively incorrect melting temperatures. The FF19SB melting temperatures appear inaccurate and imprecise as the temperature range of simulations appear too high. This instability at temperatures above 300 K is consistent with literature [11]. The observation reinforces the idea that temperature ladders that are too far above or below the melting temperatures provide insufficient information for extrapolation.

**Table 2.** *T*_*M*_, internal energy difference (ΔU) and entropy difference (ΔS) obtained for Chignolin with three force fields using interpolation. For the FF99SB force field, the reference values determined as detailed in section 3.3.2 has been included. For FF19SB, the obtained estimates of internal energy ΔU and entropy ΔS were unphysical owing to the high temperatures relative to the corresponding melting temperatures. Experimental reference values ΔH = 26.0 kJmol^−1^ and ΔS = 82.3 Jmol^−1^K^−1^.

| Force field | State | $T_M$ (K) | $\Delta U/\Delta H$ (kJ mol$^{-1}$) | $\Delta S$ (J mol$^{-1}\text{K}^{-1}$) |
| --- | --- | --- | --- | --- |
| FF99SB | Folded | 363 | 25.3 | 69.8 |
|  | Misfolded | 373 | 33.2 | 88.9 |
|  | Not Unfolded | 398 | 29.6 | 74.3 |
| FF14SB | Folded | $303 \pm 2$ | $22.3 \pm 0.9$ | $73.4 \pm 2.6$ |
| | Misfolded | $294 \pm 3$ | $22.4 \pm 1.1$ | $76.4 \pm 3.2$ |
| | Not Unfolded | $323 \pm 1$ | $22.3 \pm 0.7$ | $69.0 \pm 2.2$ |
| FF19SB | Folded | $234 \pm 47$ | - | - |
| | Misfolded | $256 \pm 44$ | - | - |
| | Not Unfolded | $260 \pm 30$ | - | - |

As a final comparison between the different force fields, the obtained internal energy and entropy differences between folded, misfolded and ‘not unfolded’ states and the unfolded state is reported (table 2). For FF99SB, the reported values were obtained from section 3.3.2. The FF19SB force field was excluded from this analysis since the obtained values are unreliable (see fig. S33 and fig. S34). For FF14SB simulations, the values shown in table 2 were obtained from interpolation. These quantities can be compared to the original results of Honda et al. [22], where the values obtained from a global fit to NMR melting curves were ΔH = 26.0 kJmol^−1^ and ΔS = 82.3 Jmol^−1^K^−1^. It should be noted that the experimental reference is the enthalpy while our simulations gave internal energy differences (owing to the NVT ensemble). Since no distinction was made between the folded and misfolded state in the experimental work, the ‘not unfolded’ state is most suited for comparison. It is easy to see that for the FF99SB force field, the internal energy difference is overestimated in comparison to the enthalpy and the entropic contribution is underestimated. However, the FF14SB force field underestimates both the internal energy contribution and the entropic contribution. The latter force field therefore displays an error correction where the consistent underestimation of both quantities leads to accurate estimates of the *T*_*M*_.

## 5 Summary and Conclusion

In this work, we examined Temperature Replica Exchange Molecular Dynamics (TREMD) simulations as a practical tool for estimating the melting temperature (*T*_*M*_) of small biomolecules. As a model system, we looked at the small peptide Chignolin. Our results confirm that TREMD is generally much more efficient than conventional MD (cMD) to study temperature-dependent equilibria of conformational states. This is in agreement with various other studies [16, 17, 32]. It has been shown that Chignolin exhibits variation in the relative stability of folded states depending on the utilized force field [10, 11]. Therefore, we determined the equilibrium between both folded states and the unfolded state across a wide range of temperatures for each force field used. We then estimated melting temperatures, Δ*U* and Δ*S* for the folded state, misfolded state, as well as the combined ‘not unfolded’ state. These quantities largely agreed with wet-lab experiments. We believe our results may be more accurate than prior computational studies (Kührová et al. [32], Okumura [33]) considering the consistency across different setups of small ladder TREMD simulations and the significantly higher simulation times over a wide temperature range. Notably, at low simulation times, our simulations reproduced the *T*_*M*_ of the ‘not unfolded’ state (fig. 9, 410-450 K below 300 ns for several starting structures) reported in these prior studies.

To formulate guidelines for the practical application of small ladder TREMD, we investigated the conditions under which these simulations are most efficient. Specifically, we considered two central aspects. Firstly, we investigated the influence of the starting conformations of replicas on the rate of convergence of TREMD. Secondly, we investigated the placement of the temperature ladders. Unsurprisingly, the optimal choice for these setup parameters was found to depend on the melting temperatures of the metastable states of the system. Generally speaking, melting temperatures may be estimated best–i.e., with high accuracy and precision, from low simulation times–if the temperature ladder overlaps with the melting temperature(s) of interest. The choice of the starting conformations also has a strong impact on rate of convergence of the simulations. For our example, convergence of TREMD simulations was observed to be considerably slower when comparing the worst and best possible setups, with the worst-case taking more than five times longer to equilibrate (seen in fig. 10 and figs. S9 to S13).

Our simulations point to an iterative scheme as a solution to this mutual dependence of melting temperatures and setups. Small temperature ladders may first be simulated at high temperatures and then progressively shifted down with more informed choices of starting structures. While small ladders, particularly at relatively low temperatures can suppress the enhancement of sampling, it was seen from the FF99SB simulations that low temperature simulations with an appropriate set of starting structures can provide reasonable first estimates of the melting temperatures. When no information regarding state probabilities is available, a good first choice would be high-temperature simulations starting from unfolded structures. Subsequent simulations can then be planned such that the temperature ladders overlap with the *T*_*M*_ estimates from earlier runs and the fraction of replicas initialized with each metastable state matches the mean probability of that state across the ladder. In essence, we propose a separate construction of small segments of the melting curve, which could potentially be more resource-efficient than using a single large temperature ladder (depending on hardware).

Prior work suggested that the equilibrium distribution of replicas among metastable states corresponds to the mean state probability over the temperature ladder. This result is also validated here. While largely intuitive, the results further showed that modest deviations in the number of replicas per state with respect to the equilibrium distribution are acceptable for starting structures. However, it was also noted that a mixture of multiple starting structures generally outperforms a single starting conformation. This was more so for short simulations (in time). This behavior is consistent with the predictions of the OU model discussed in section 2.4.

The notion of a small ladder is inherently system dependent. As the system grows large, more replicas might be necessary to cover the temperature range of interest. Similarly, the temperature range spanned by a certain replica count shrinks as the system increases in size and complexity. For such systems, in which a single temperature ladder may not be able to span the temperature range of interest, the proposed scheme is a promising solution.

## Acronyms

T_M_: melting temperature
cMD: conventional MD
IQR: inter-quartile range
MCMC: Markov Chain Monte Carlo
MD: Molecular Dynamics
MDW: Model Double-Well
MFPT: mean first passage time
MSM: Markov state model
OU: Ornstein-Uhlenbeck
PT: Parallel Tempering
TREMD: Temperature Replica Exchange Molecular Dynamics

## Author contributions statement

N.K.R performed simulations, wrote programs and scripts, analyzed data, designed mathematical model, wrote paper. P.K.Q. designed project, analyzed data, wrote paper, analyzed mathematical model, M.Z. designed project, supervised computational work, provided computational resources, wrote final manuscript version. N.K.R., P.K.Q. and M.Z. analyzed results, reviewed the manuscript.

## Data Availability

Relevant code and data will be made available upon final publication.

## Acknowledgments

The authors thank Martin Kulke for useful discussions. This work was financially supported by the Deutsche Forschungsgemeinschaft (DFG Za153/29-1) and by local compute cluster (partially funded by DFG INST 95/1610-1 FUGG). We acknowledge additional HPC resources provided by the Erlangen National High Performance Center (NHR@FAU).

## A Calculation of the Ornstein-Uhlenbeck Error

The Gaussian noise approximation can be understood as follows. For a two-state system, each frame in a trajectory is essentially the toss of a biased coin. The probability of F is *p*_*F*_ and the probability of U is *p*_*U*_ = 1 − *p*_*F*_. If there are Δ*t* frames in total, the probability with which Δ*t*^*′*^ frames are in F and Δ*t* − Δ*t*^*′*^ frames are in U is given by 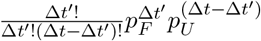. As the total number of frames in each block increases Δ*t*, the probability mass described just now tends towards a Gaussian function with mean Δ*tp*_*F*_ and variance Δ*tp*_*F*_ *p*_*U*_. Hence, it can be assumed that typically, the fluctuations about the true probability for large enough blocks would be Gaussian distributed. However, initially, the set of starting structures skews the values sampled from this Gaussian distribution. The presence of one state in more replicas leads to an increased sampling of the tail end of the Gaussian distribution of block probabilities. The variance of the estimator is therefore relatively high for short time scales, which is what the deterministic initial conditions encode in the Ornstein-Uhlenbeck (OU) model.

The above picture also assists in understanding how the two-state model considered here extends to multi-state systems. Since every multi-state system can be reduced to a set of two-state systems (reducing it to state A or not in state A for every A), one can apply the same arguments as above, although with different probabilities.

### A.1 Variance of the OU Model

To determine the uncertainty analytically, it is necessary to establish a preliminary result. Namely, it is necessary to find 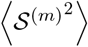 as regardless of whether bootstrap or jackknife is used to estimate the standard error of the mean, the quantity being summed is the same.

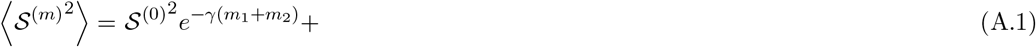

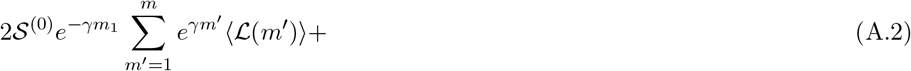

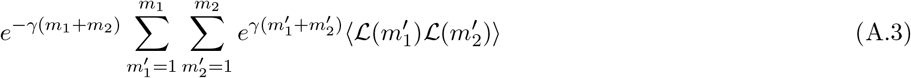

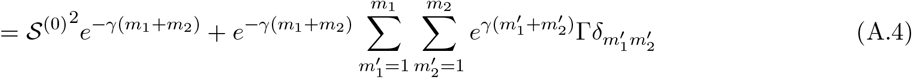

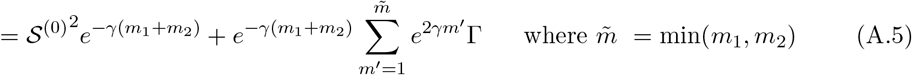

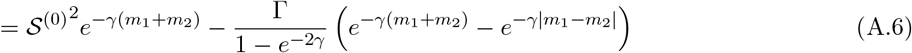

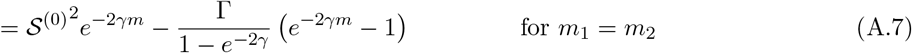

In our calculations, we can set

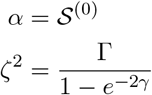

### A.2 Uncertainty in Metastable-State Probability Estimates via Bootstrapping

The bootstrap-based formulation of the variance of the estimator of *s* is straightforward to calculate given the OU model variance. Consider the blocked average of the state function 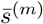 at block index *m*. Assume that there are *M* blocks in total. Now consider *B* bootstraps of these blocks, choosing *n*_*r*_ blocks at random each time so that the statistics for a simulation time of *n*_*r*_Δ*t* can be extracted. Suppose that for bootstrap index *b*, one has *n*_*r*_ blocks sampled which are indexed as *m*_*b,j*_. For such as a bootstrap, the variance of the estimator can be expressed as follows

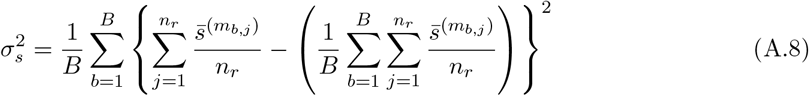

Here, we may assume that the latter term, namely

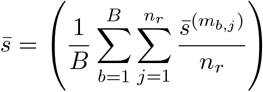

tends towards the mean 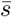 as the number of bootstraps increases. This implies that the terms in the variance can be reformulated as

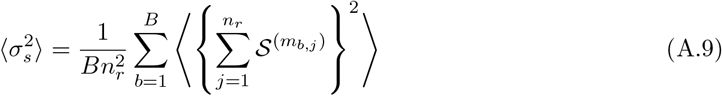

The square of a sum is easily expanded and the ensemble average may be spread over each term. This gives

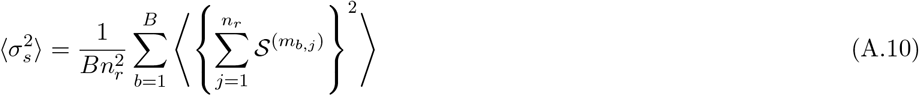

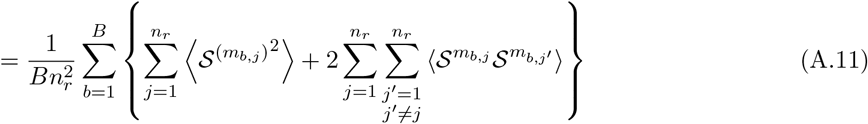

This expression, using eq. (A.7) and prior steps, gives the following result:

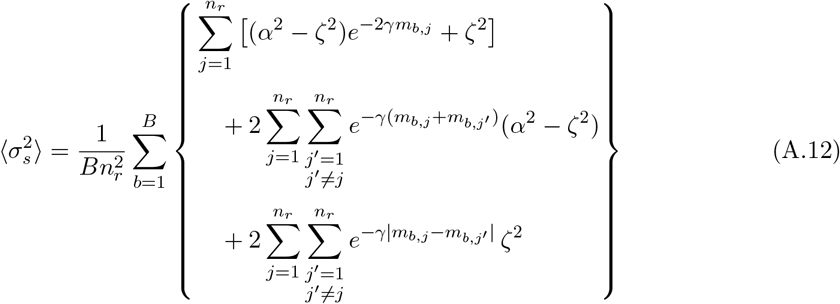

If one considers a large number of bootstraps from the blocked means of the state function 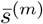, then it is good to assume that all blocks are equally likely to be chosen. This means that the quantities within curly braces above can be replaced with a sum over sequential numbers *m* = 0, … *N* (*N* being different values for each sum) and weighted by a probability mass dependent on *m*. To make this more concrete, the following can be said. Given a sequence of numbers 1, …, *M*, the probability of choosing two of them without replacement such that their sum is equal to *m* is given by

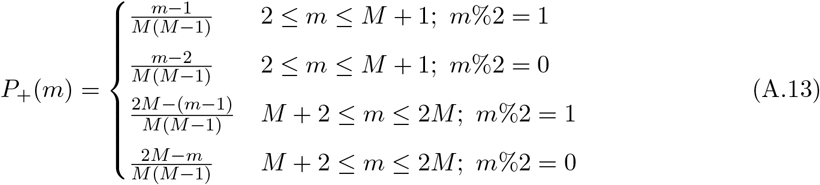

and choosing them without replacement such that the the difference is *m* is given by

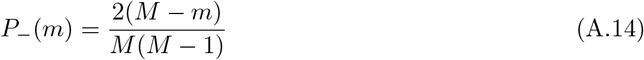

Hence, it is straightforward to calculate each quantity in eq. (A.12) above.

1. The first term is a sum over a single block index *m*_*b,j*_. For a large number of bootstraps, each value is going to be sampled approximately 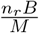 times (the probability of sampling each point is 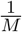 and there are *n*_*r*_*B* samples in total). This means that the sum of *B* and *n*_*r*_ can be replaced by a sum over all terms multiplied by 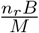.

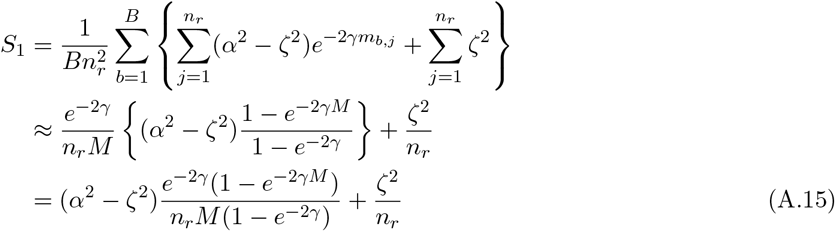
2. The next term is a sum of two block indices. This term can be dealt with using eq. (A.13). Once more, we sample *n*_*r*_(*n*_*r*_ − 1)*B* pair values in total and the probability of each value is as given in expression eq. (A.13). The expression can be split into four sums based on whether *m > M* + 1 and on the parity of *m*. Using this simplification and the arithmetico-geometric series sum we have

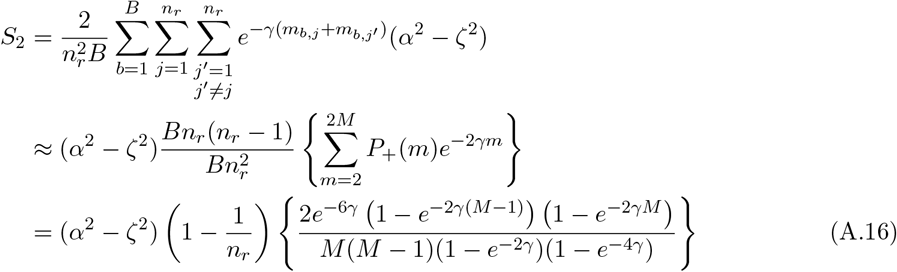
3. The third and final term is the sum over the absolute difference of indices. This is once more modeled using eq. (A.14) and the total pair count *n*_*r*_(*n*_*r*_ − 1)*B*. Here, once more we perform a series sum

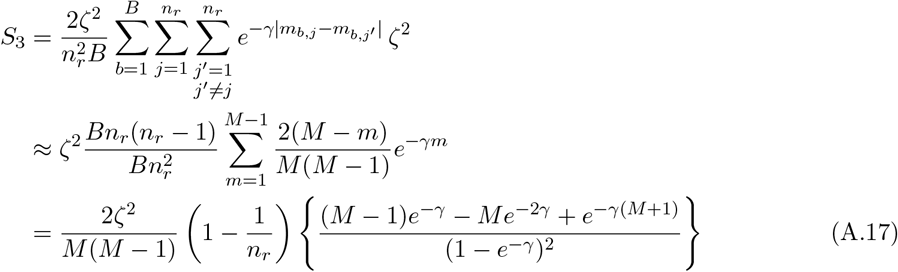

Putting eq. (A.15), eq. (A.16), and eq. (A.17) together, we get

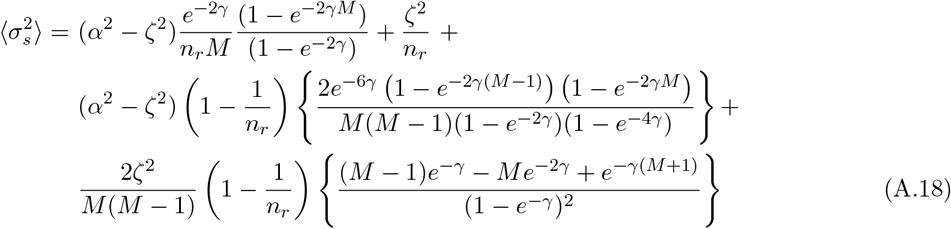

Within this framework, there are the known contributions to the uncertainty from jackknife. At the same time, there are two new terms that appear in the expression. Now, consider the limit of small *γ*, small *M* and small *γM* i.e. long relaxation times and short simulations. We have

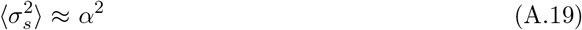

However, if *M* is fairly large but still small compared to *γ*^−1^, the expression also reduces to

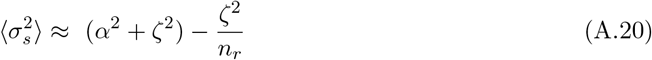

Here, it is increasing because the variance is underestimated for short simulations. As the estimate of the mean approaches the true mean, this expression plateaus and then decreases. On the other hand, for long simulation times or large *M* and large *γM*, we have

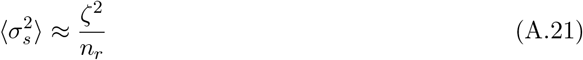

since all terms inversely dependent on *M* become negligible. While this is the limit of infinite simulation time, realistic simulations would have the intermediate limit with large *M* and *γM* = *u* ∼ 𝒪(1). Finally, it can be seen from the expression in eq. (A.18) that the terms with *α* and *ζ* can be collected to give

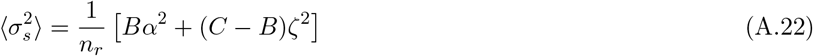

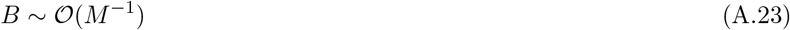

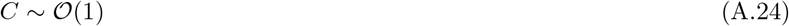

which is the exact same form as the jack-knife error owing to *n*_*r*_ ∈ [1, *M*]. The effect of initial conditions scales as *M*^−1^ and the overall uncertainty decays as 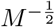.

