## Supplemental Information for "Estimation of Protein Melting Temperatures Using Small-Ladder Replica Exchange Simulations"

#### Acronyms

$T_M$ melting temperature

**cMD** conventional MD

**MCMC** Markov Chain Monte Carlo

**MDW** Model Double-Well

**PT** Parallel Tempering

**TREMD** Temperature Replica Exchange Molecular Dynamics

### S1 Supplementary Material for Mathematical Model

#### S1.1 Note on the Energy Surface for MCMC

The projection is assumed to be that of the free energy, i.e. a PMF and not the potential for a simple reason. Assume that $U(\vec{x}^N)$ represents the potential energy of the molecule in exact phase space coordinates $\vec{x}^N$. Assume that the free energy $F$ is projected onto some variable $\Xi \equiv \phi(\vec{x}^N) \in \mathbb{R}$. Then, one has the following definition of the free energy in terms of the potential as detailed in Hartmann et al. [1]

$$F(\xi) = -k_B T \ln \int e^{-\beta U(\vec{x}^N)} \delta\left(\phi(\vec{x}^N) - \xi\right) d\xi = U(\xi) - k_B T \ln \Omega(\xi) \tag{S1.1}$$

which is effectively a representation of the fact that the probability of a state within the range $[\xi, \xi + d\xi)$ contains all $\vec{x}^N$ for which $\Xi = \xi$. If one were to perform a Metropolis Hastings Markov Chain Monte Carlo (MH-MCMC) simulation in the projection of the internal energy, the phase space volume corresponding to each value of the collective variable would not be captured properly. Another perspective would be that according to the Boltzmann definition, the entropy is given by $S = k_B \ln \Omega$ where $\Omega$ is the density of states. If one performs a simulation of the projection of the internal energy, each value of the collective variable would have probability

$$P(\xi) = \frac{e^{-\beta U(\xi)}}{Z} \tag{S1.2}$$

which leaves out a measure of the phase space volume that actually corresponds to the single value of the collective variable. On the other hand, performing MH-MCMC simulations in the space of the free energy lends the following probability to each value of the collective variable

$$P(\xi) = \frac{e^{-\beta F(\xi)}}{Z} = \frac{e^{-\beta[U(\xi) - k_B T \ln \Omega(\xi)]}}{Z} = \frac{\Omega(\xi) e^{-\beta U(\xi)}}{Z} = P(\{\vec{x}^N : \phi(\vec{x}^N) \in [\xi, \xi + d\xi)\}) \tag{S1.3}$$

which is weighted with the density of states.

#### S1.2 Formulation of the Ornstein-Uhlenbeck Model Using Jackknife

The jackknife error can be used for calculating the error variation as well. Consider the jackknife error

$$\sigma_s = \sqrt{\frac{M-1}{M}} \sqrt{\sum_{m=1}^{M} (\bar{s}^{(m)\prime} - \bar{\bar{s}})^2} \tag{S1.4}$$

$$\bar{s}^{(m)\prime} = \frac{1}{M-1} \left\{ \sum_{i=1}^{M} \bar{s}^{(i)} - \bar{s}^{(m)} \right\} = \frac{1}{M-1} \left\{ M\bar{s} - \bar{s}^{(m)} \right\} \tag{S1.5}$$

$$\bar{\bar{s}} = \frac{1}{M} \sum_{m=1}^{M} \bar{s}^{(m)\prime} = \frac{1}{M(M-1)} \sum_{m=1}^{M} \left( M\bar{s} - \bar{s}^{(m)} \right) = \bar{s} \tag{S1.6}$$

$$\begin{aligned} \implies \sigma_s &= \sqrt{\frac{M-1}{M}} \sqrt{\sum_{m=1}^{M} \left( \frac{(M-1)\bar{s} - M\bar{s} + \bar{s}^{(m)}}{M-1} \right)^2} \\ &= \sqrt{\frac{1}{M(M-1)}} \sqrt{\sum_{m=1}^{M} \mathcal{S}^{(m)2}} \end{aligned} \tag{S1.7}$$

From here, we simply replace $\mathcal{S}^{(m)}$ with Equation A.7. This gives

$$\sqrt{\langle \sigma_s^2 \rangle} = \sqrt{\frac{1}{M(M-1)}} \sqrt{\sum_{m=1}^{M} \left\{ \mathcal{S}^{(0)2} e^{-2\gamma m} + \frac{\Gamma}{1-e^{-2\gamma}} \left(1 - e^{-2\gamma m}\right) \right\}} \tag{S1.8}$$

$$= \sqrt{\frac{\Delta t}{t_{\mathrm{sim}}(t_{\mathrm{sim}} - \Delta t)}} \sqrt{\left( \mathcal{S}^{(0)2} - \frac{\Gamma}{1-e^{-2\gamma}} \right) e^{-2\gamma} \frac{1 - e^{-2\gamma \frac{t_{\mathrm{sim}}}{\Delta t}}}{1 - e^{-2\gamma}} \Delta t + \frac{\Gamma}{1 - e^{-2\gamma}} t_{\mathrm{sim}}} \tag{S1.9}$$

### S2 Supplementary Material for Materials and Methods

#### S2.1 Model Double-Well (MDW) Construction

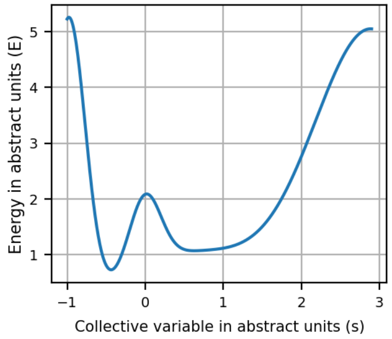

**Fig. S1:** The MDW potential used for performing Markov Chain Monte Carlo simulations. The folded state minimum can be seen on the left and the unfolded state on the right. The width is meant to be proportional to the entropy of the metastable state. As one might expect and also based on fig. S3, the unfolded state is more flexible than the folded state. The barrier is at $s = 0.015$.

| Scaling factor | Mean | Std. deviation |
|---|---|---|
| 5.00 | -0.98 | 0.21 |
| -0.15 | -0.51 | 0.20 |
| 1.24 | 0.00 | 0.20 |
| 5.00 | 2.90 | 0.75 |
| 1.01 | 0.57 | 0.95 |

The MDW was constructed using a sum of Gaussian functions. The mean, variance and scaling factor of each Gaussian has been provided below

Walls were also defined for the MDW at $s = -0.98$ and $s = 2.9$ with infinitely high walls. This effectively meant that the particle would be reflected from the walls. Simulations were performed at the following beta values

$$\beta = 13.45, 12.08, 10.87, 9.81, 8.86, 8.03$$

These values were not chosen to optimize the exchange success rate, which was close to 80 %. It is known that an extremely high exchange success rate leads to over-mixing and thereby diminishing the benefits of the parallel tempering steps. However, simulations were performed with the following parameters

```
        n_ex       = 10000      # Number of exchanges
        n_steps    = 25000      # Number of steps between exchanges
        nwx        = 100        # Trajectory written every 100 steps
        step_size  = 0.05       # step_size
        prf        = uniform    # Uniform proposal function
```

Additionally, the starting positions were adjusted to mimic different sets of starting structures. The simulation indices and their starting structures have been provided below

**Table 1:** The setups and repetitions of Markov Chain Monte Carlo (MCMC) simulations. Replicas starting from the 'Folded' state were initialized at point -0.5. Replicas starting from the 'Unfolded' state were initialized at point 0.75.

| **Setup Idx** | **Simulation type** | **Simulation indices** | **#Folded** | **#Unfolded** |
|---|---|---|---|---|
| 1 | Conventional | 1-5 | 0 | 6 |
| 2 | Parallel tempering | 6-10 | 0 | 6 |
| 3 | Parallel tempering | 11-15 | 1 | 5 |
| 4 | Parallel tempering | 16-20 | 2 | 4 |
| 5 | Parallel tempering | 21-25 | 3 | 3 |
| 6 | Parallel tempering | 26-30 | 4 | 2 |
| 7 | Parallel tempering | 31-35 | 5 | 1 |
| 8 | Parallel tempering | 36-40 | 6 | 0 |

#### S2.2 Chignolin

The stable states of Chignolin are shown in fig. S2.

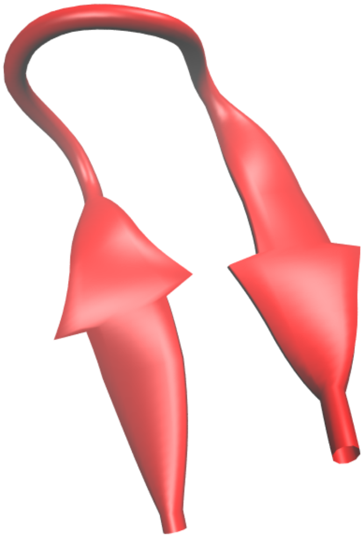

**(a)** Folded

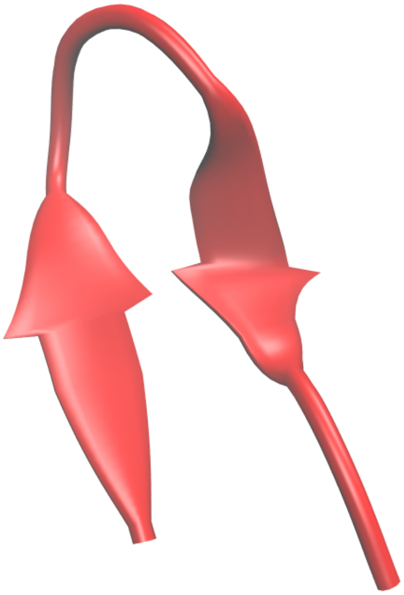

**(b)** Misfolded

**Fig. S2:** Shows the folded and misfolded states of Chignolin obtained with the FF99SB force field. They were used as references for all subsequent simulations. While several sub-states were observed, they were largely fluctuations of these two metastable states and therefore dealt with as such. Chignolin was treated as a three-state system with a folded, misfolded and unfolded state.

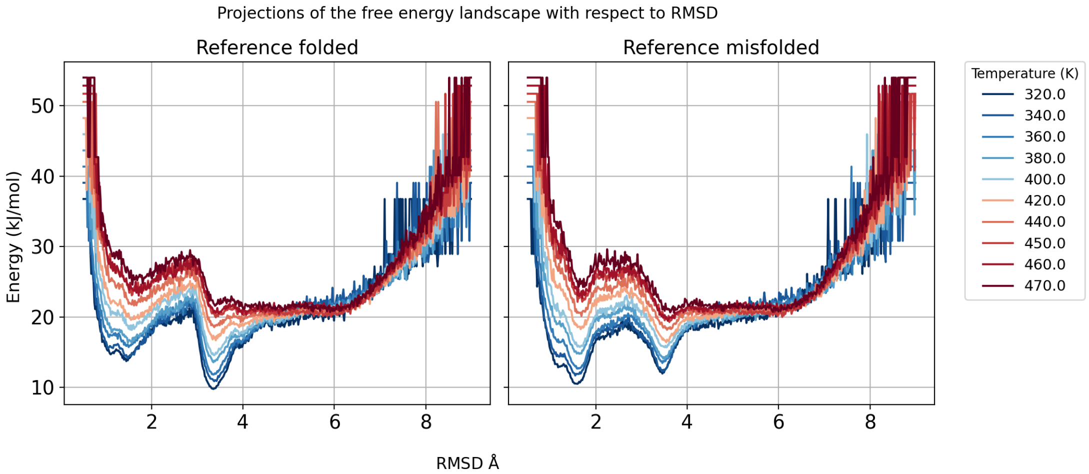

**Fig. S3:** The projection of the free energy distribution on the RMSD with respected to the native and non-native folded states of Chignolin with the FF99SB force field.

##### S2.3 Different Resampling Schemes

###### S2.3.1 ${}^{m}C_m$ - bootstrapping

For block counts $m = 1, \ldots, M$, $b$ bootstrap resamples were drawn from the first $m$ blocks. The resulting bootstrap variance ($\langle \sigma_s^2 \rangle$) provided an estimate of the standard error ($\sqrt{\langle \sigma_s^2 \rangle}$) of state probabilities for simulations truncated after $m$ blocks. As $m$ increased, the variance initially grew, then plateaued, and eventually decayed approximately as $1/m$ for all setups. The initial underestimation of variance was expected, as the influence of the starting structures gives precise but biased estimates. Multiplying the variance by $m$ yielded coefficients that either increased or decayed toward a constant, depending on the starting structures. Let $k$ denote the number of equilibration blocks.

###### S2.3.2 ${}^{M}C_m$ - sub-sampling with replacement from all blocks

From the full set of $M$ blocks, $m = 1, \ldots, M$ blocks were resampled (with replacement) $b$ times. The variance of the resampled state-probability estimates decayed as $1/m$. Scaling the variance sequences with $m$ returned constants of proportionality for each state and replica.

###### S2.3.3 ${}^{M-k}C_m$ - sub-sampling with replacement from equilibrated blocks

Resampling $m = 1, \ldots, M - k$ blocks from the $M - k$ blocks following the first $k$ blocks gave $1/m$ decay as well. However, the constants of proportionality differed when resampling with and without the first $k$ blocks.

#### S2.4 Reference melting temperature ($T_M$) Calculation for cMD

The reference melting temperatures obtained for Chignolin with FF99SB force field and TIP3P water model at 0.23 M salt concentration from conventional MD (cMD) have been provided here. The two approaches for $T_M$ estimation are the van't Hoff analysis and softmax fitting. Roughly 193 $\mu$s of cMD simulation time split over 9 temperature points were performed. The van't Hoff curve for these has been provided in fig. S4.

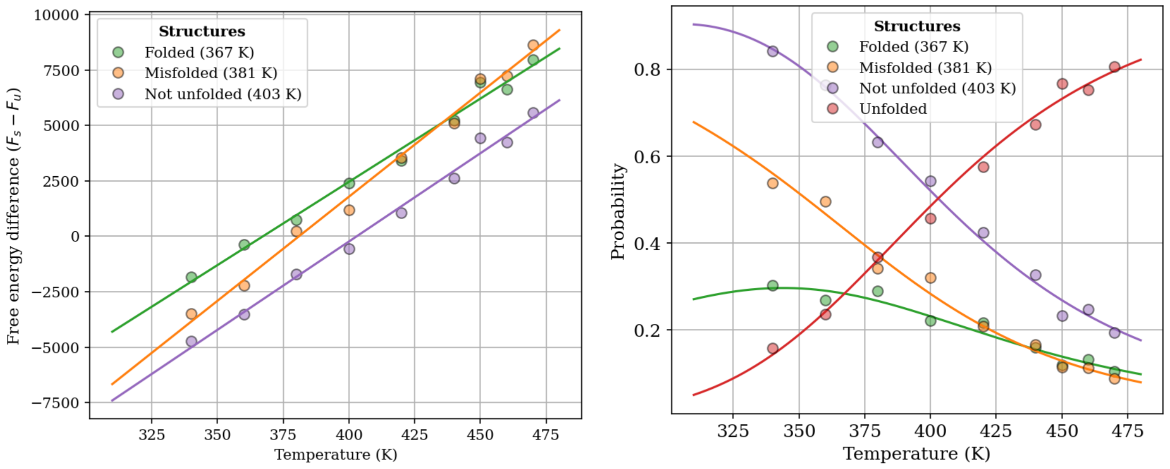

**(a)** Linear fits

**(b)** Softmax inversion of linear fits

**Fig. S4:** Using a van't Hoff analysis of the probability estimates made using the full simulation from conventional MD simulations, the folded, misfolded and combined states had $T_M$ estimates of 367 K, 381 K and 403 K respectively. The free energy estimates obtained by inverting the probabilities as well as the linear fits to these points have been shown here.

Similarly, unconstrained sigmoidal fits were also performed to this data to generate melting curves. These results are shown in fig. S5.

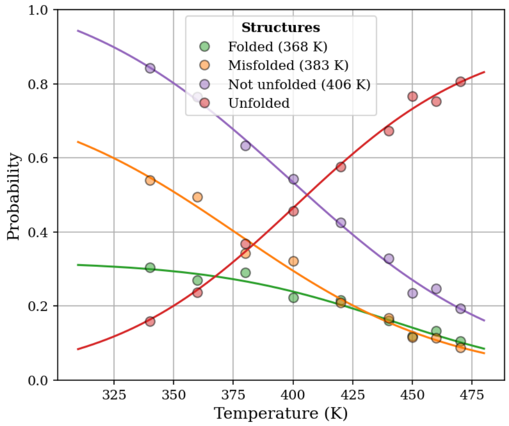

**Fig. S5:** Unconstrained fits to the probability distributions obtained from cMD simulations of Chignolin. Unconstrained fits are fast and stable but as should be clear from the figure, extrapolating this distribution to temperatures beyond the range of the simulations would yield unphysical results. The sum of probabilities would exceed one. This can be mitigated by performing constrained fits to each melting curve such that their sum is one. However, constrained fits are unstable. This effectively renders this method unusable for the short-ladder scheme prescribed in this study.

While cMD simulations generated probability distributions that appeared fairly well converged, we also leveraged the vast quantities of Temperature Replica Exchange Molecular Dynamics (TREMD) simulations performed with different starting structures. Using fig. 6, it was easy to see that most simulations yielded the same distribution of replica count per metastable state after an initial equilibration period. The full version of fig. 6 is give in figs. S14 and S15. Equilibration phases were removed and the data from equilibrated TREMD simulations were pooled to generate reference values. The data for the TREMD reference has been provided in the main text (section 3.3.2).

In comparison to the latter set of reference values, the cMD simulations appear far from convergence, with melting temperatures being 5-10 K from the values in fig. 4.

### S3 Supplementary Material for Results and Discussion

A setup in this context refers to the combination of starting structures and temperature ladder.

#### S3.1 Minimal Model

The $T_M$ estimates obtained from conventional and Parallel Tempering (PT)-Markov Chain Monte Carlo (MCMC) have been provided in fig. S6 for three cases – sub-sampling with replacement from full simulations, bootstrapping for increasing block counts, and sub-sampling with replacement from equilibrated simulations.

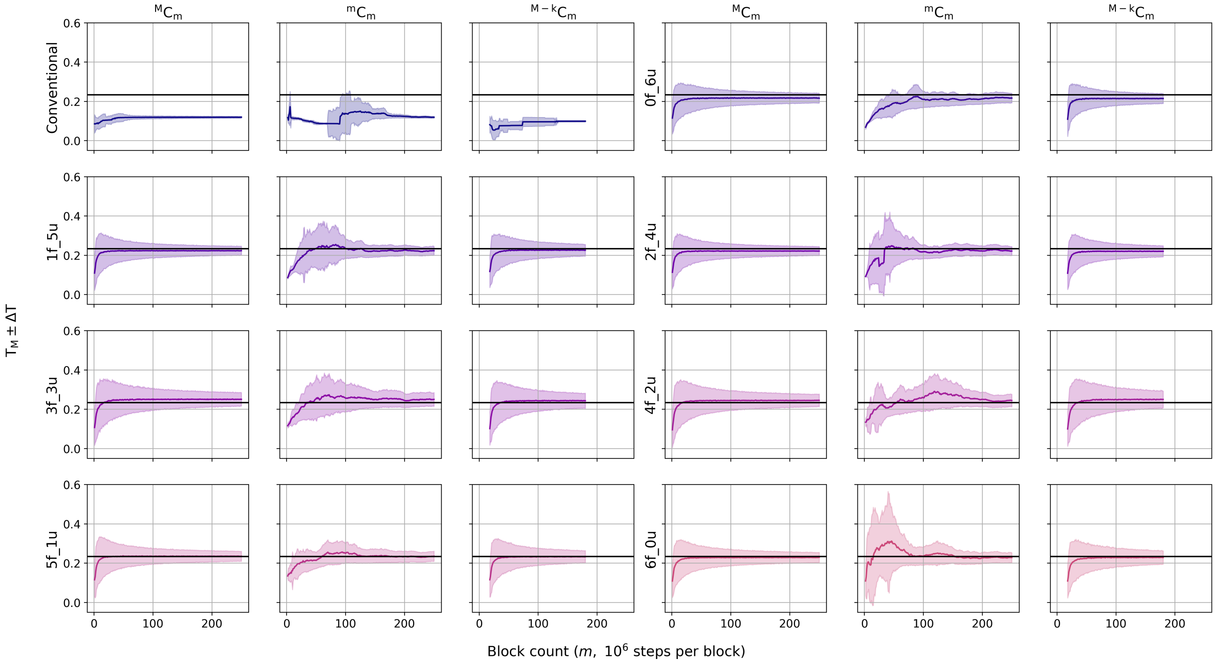

**Fig. S6:** The $T_M$ estimates from MCMC simulations averaged over 5 repetitions for each setup. The expected value of the $T_M$ for the system modeled by the MDW is 0.234 units and is shown by the solid black line. The estimates were obtained by calculating the bootstrapped mean and standard deviation and then averaging over the five iterations of the setup. Block sizes were chosen to be $10^6$ steps.

A comparison between the constants of proportionality between sub-sampled standard error estimates from $M$ blocks and $M - k$ blocks (with and without equilibration phase) has been provided in fig. S7.

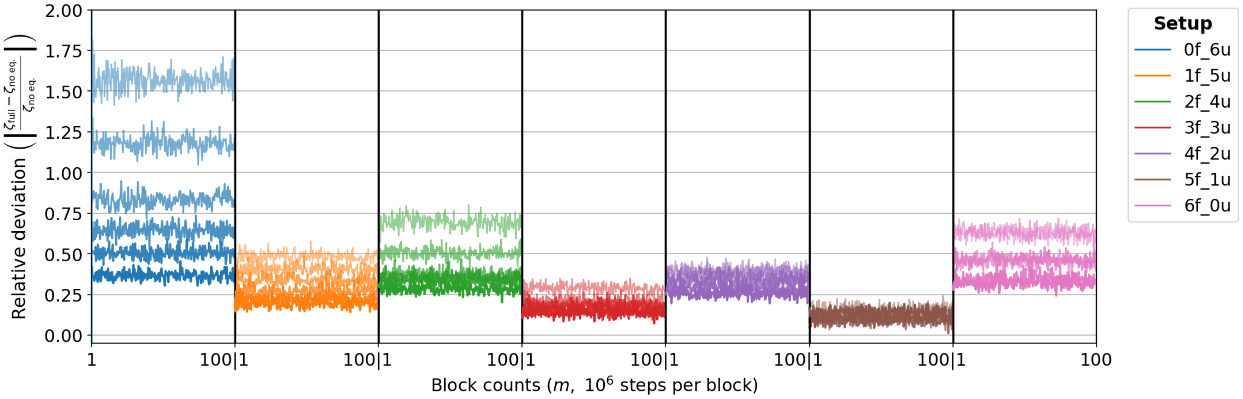

**Fig. S7:** $\zeta_{\mathrm{full}}$ and $\zeta_{\mathrm{no\ eq.}}$ are the constants of proportionality of the resampled standard error estimate with and without the equilibration phase. The plot shows $\frac{\zeta_{\mathrm{full}}-\zeta_{\mathrm{noeq.}}}{\zeta_{\mathrm{full}}}$.

The constants of proportionality $\zeta$ for the full simulation and the simulation without the equilibration phases obtained by multiplying the time series of error by the block count have been provided in fig. S8.

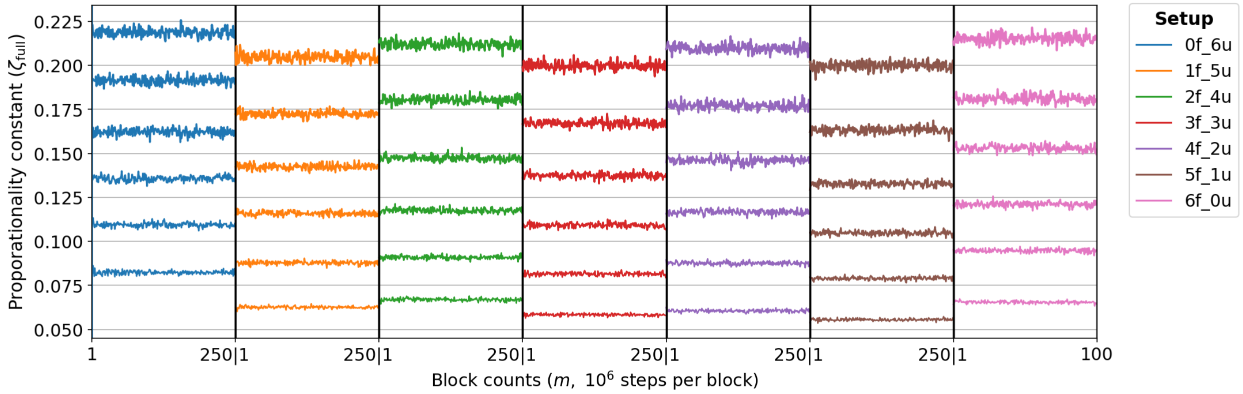

**(a)** Resample with replacement from full simulation

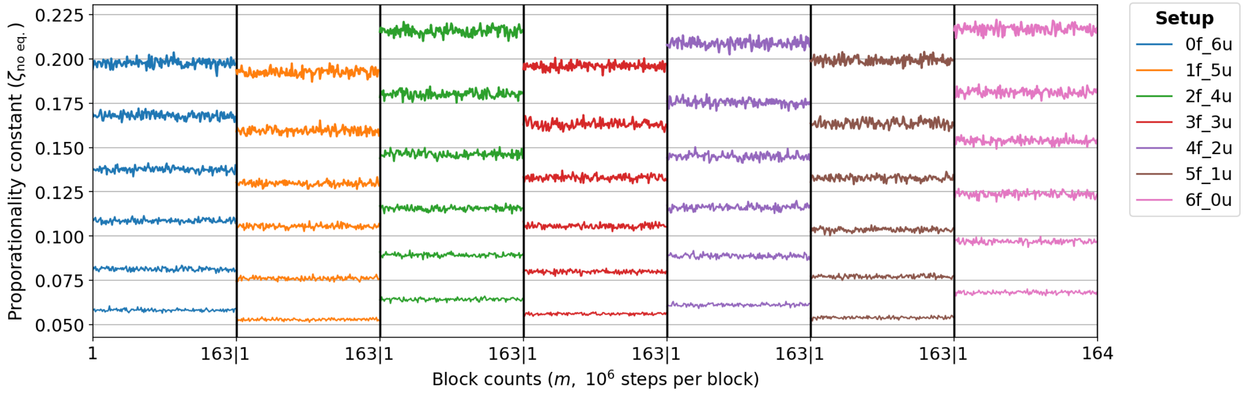

**(b)** Resample with replacement excluding equilibration phase

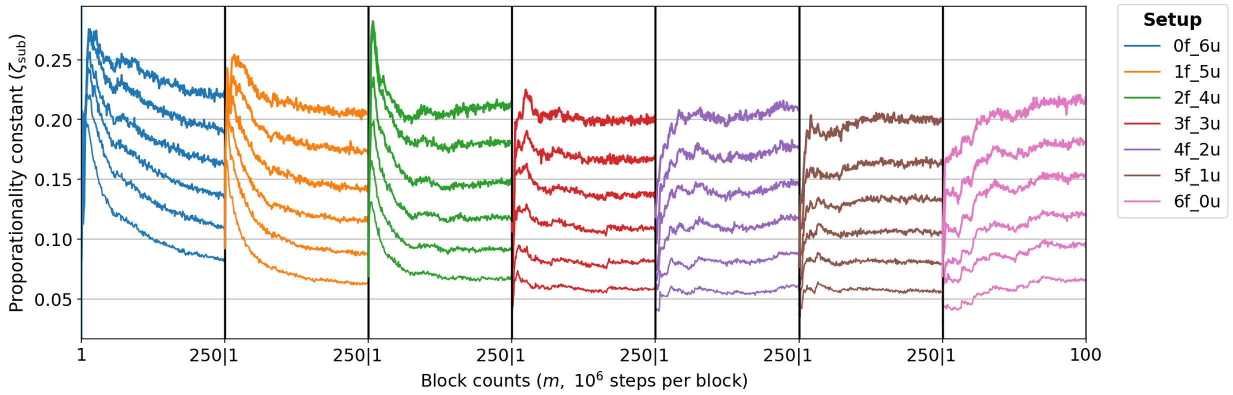

**(c)** Bootstraps

**Fig. S8:** Proportionality constant $\zeta$. Figure S8a shows results obtained by resampling from all $M$ blocks, while fig. S8b shows results obtained after excluding the first 35 % of blocks prior to resampling. Figure S8c shows the bootstrapped errors for the MCMC simulations. Angular brackets denote ensemble averaging over 5 repetitions of each setup. The resulting constants of proportionality for the standard error are denoted by $\zeta$. Line thickness is proportional to temperature. Each column therefore corresponds to a distinct setup, and each line to the $\zeta$ for an individual replica. A systematic shift in $\zeta$ is observed between the two cases (resample from $M$ and $M - k$). Setups showing the smallest change on exclusion of the equilibration blocks are those initialized with a high proportion of the folded state.

#### S3.2 Small Ladder TREMD for Melting Temperature Estimation

The bootstrapped data was used to obtain statistics for $T_M$ estimates. Please refer to fig. 3 and table 1 for definition of temperature ladders and setups. Ladder one was divided into 15 blocks, while all other ladders were divided into 30 blocks.

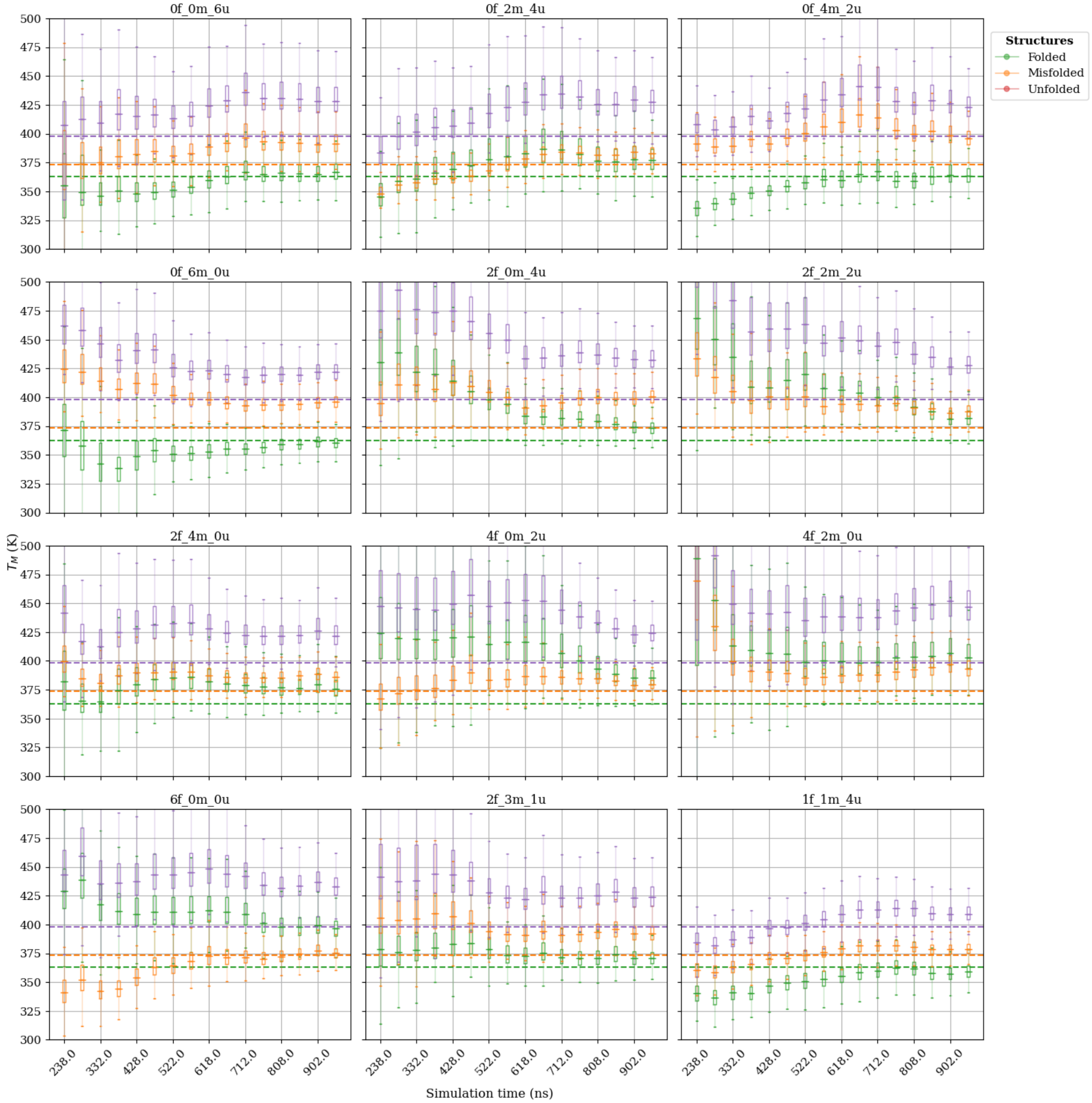

**Fig. S9:** Melting temperatures obtained by extrapolating ladder 1 simulations.

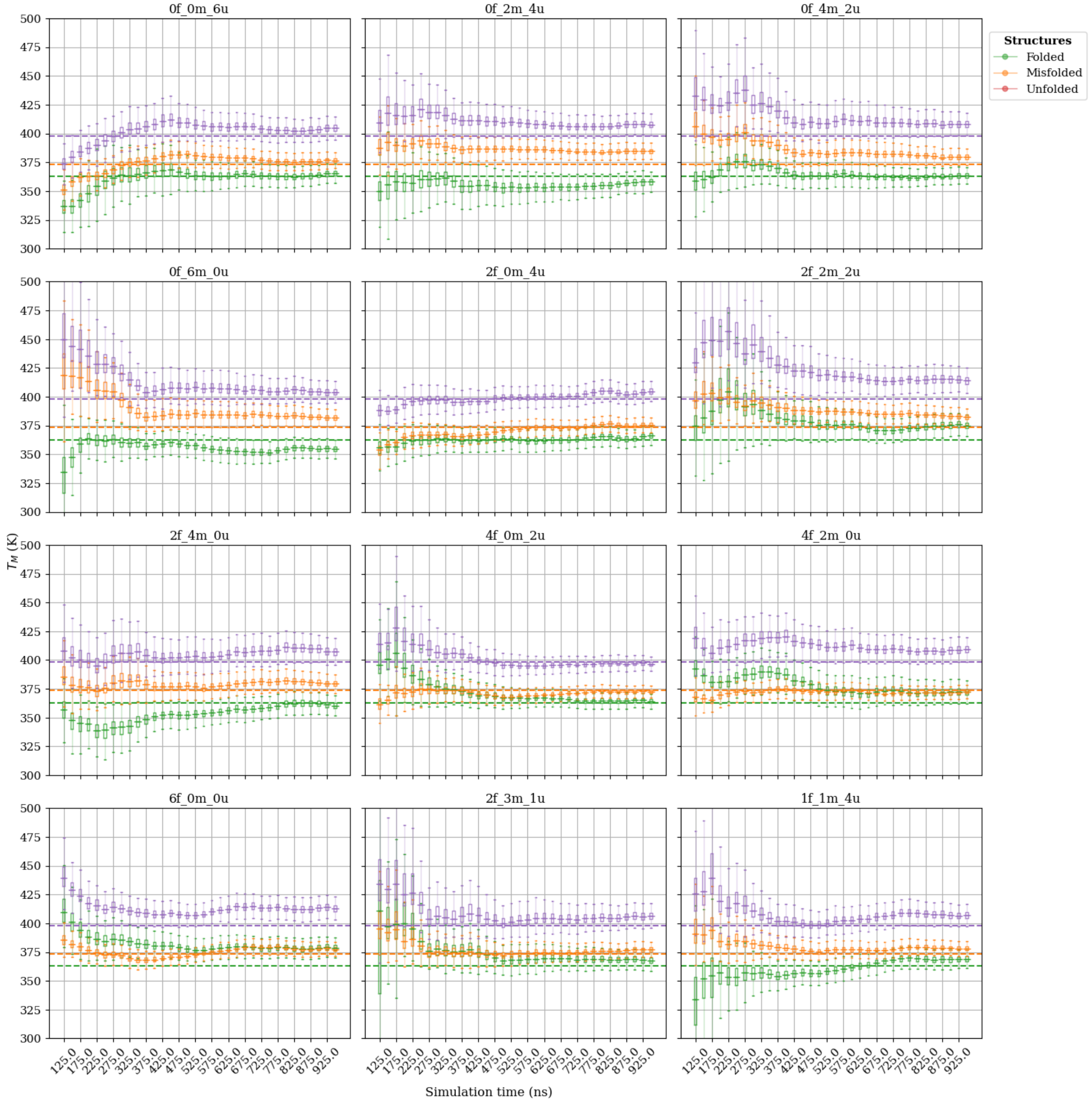

**Fig. S10:** Melting temperatures obtained by extrapolating ladder 2 simulations.

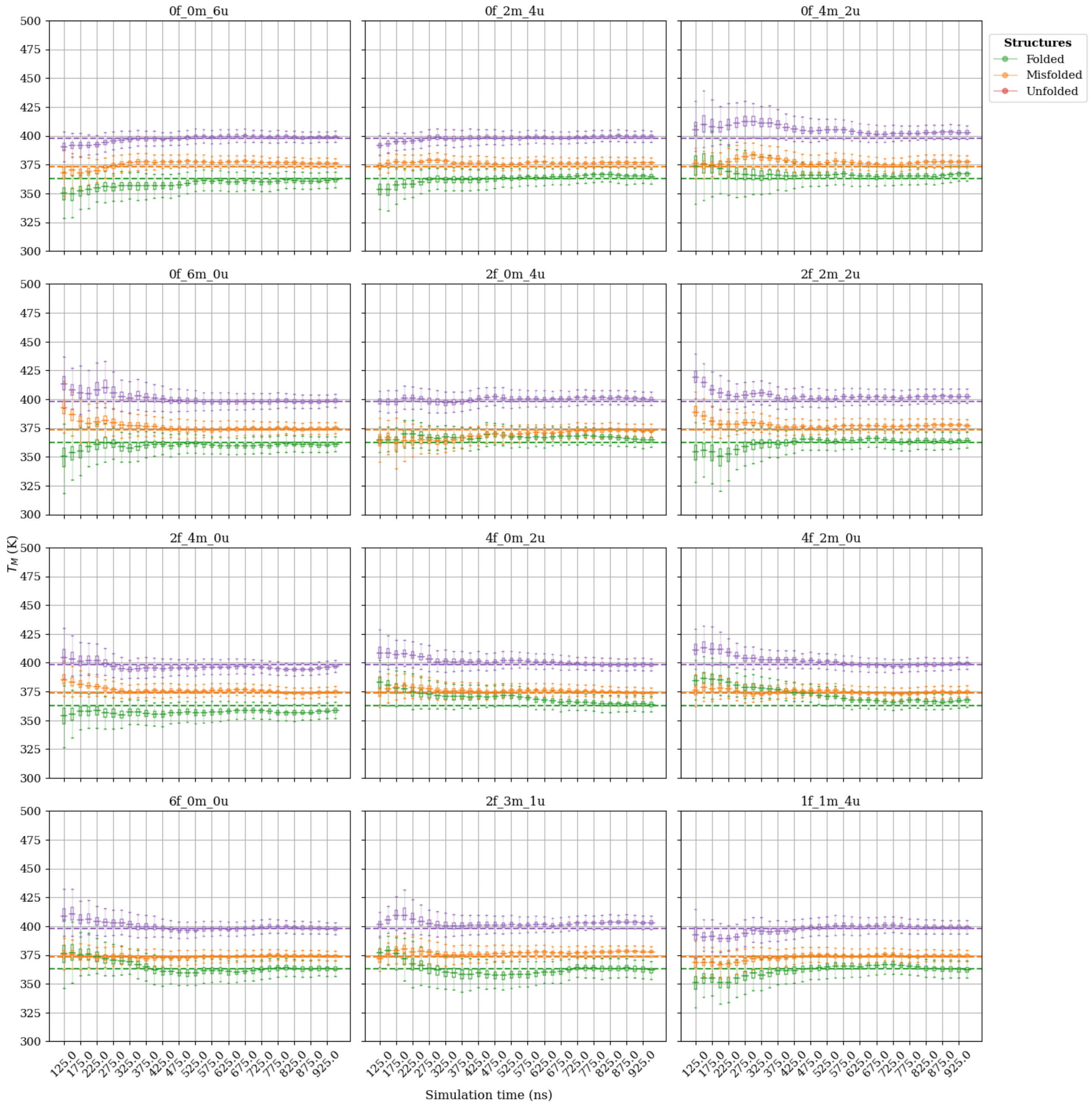

**Fig. S11:** Melting temperatures obtained by extrapolating ladder 3 simulations.

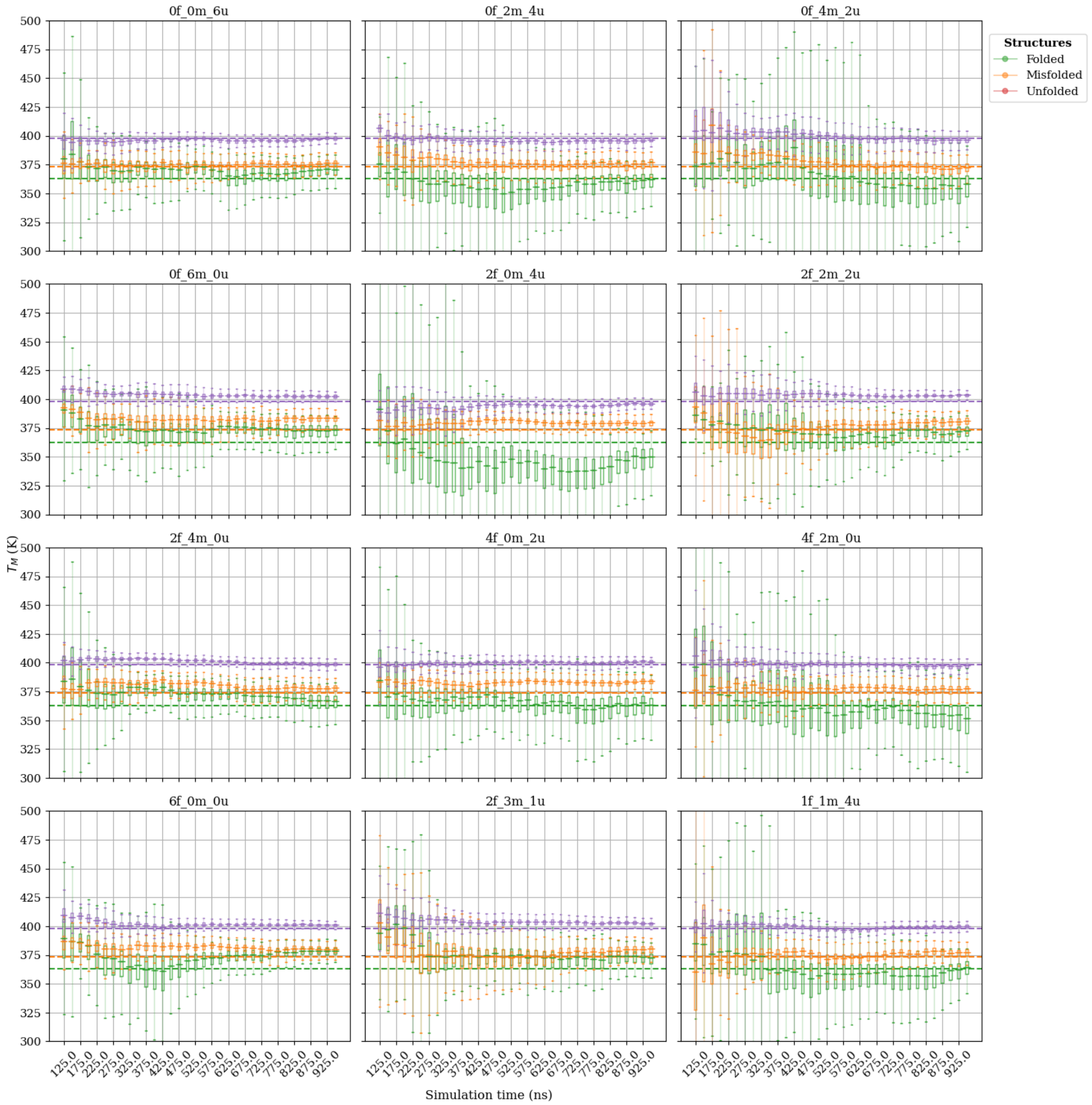

**Fig. S12:** Melting temperatures obtained by extrapolating ladder 4 simulations.

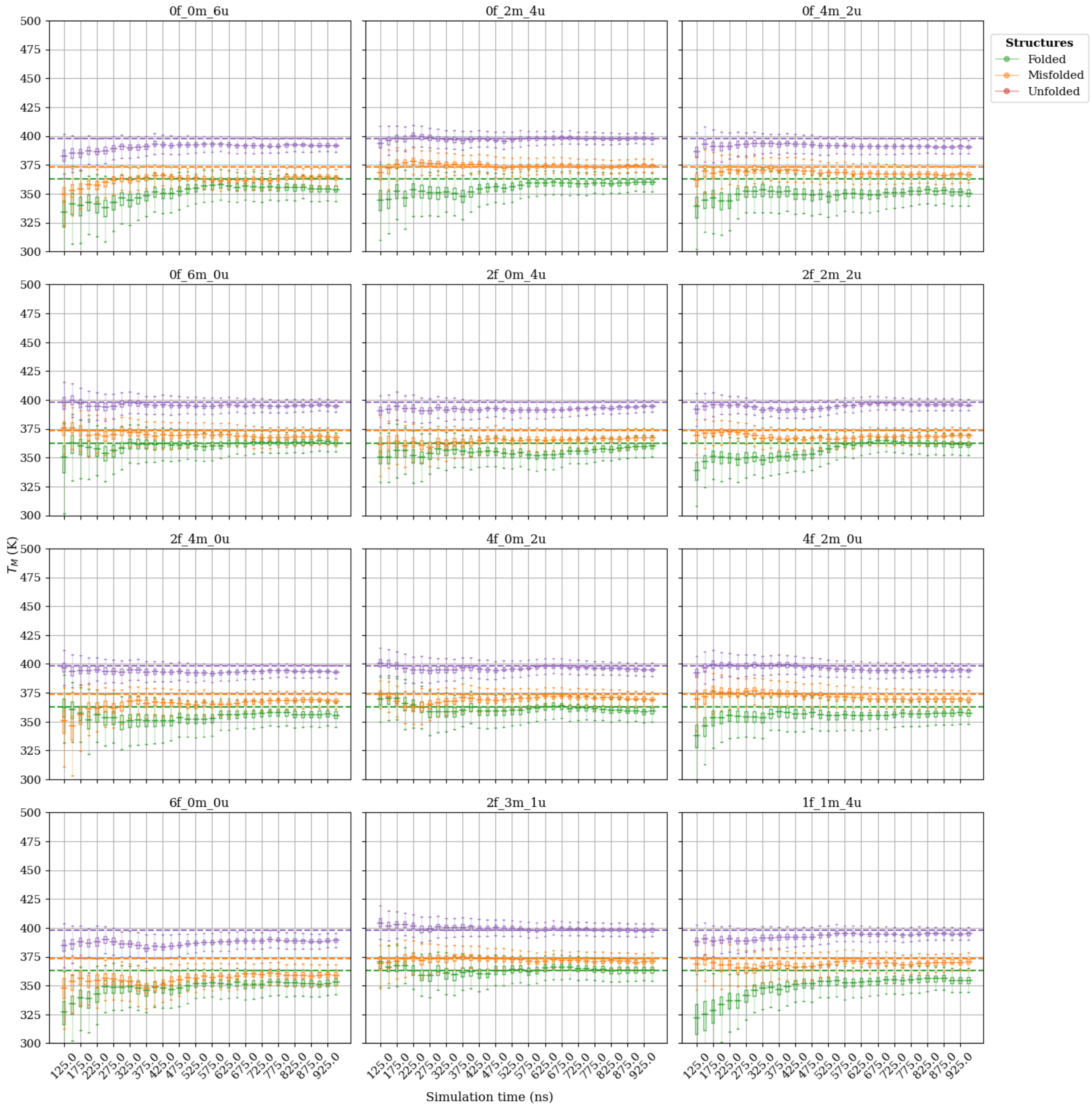

**Fig. S13:** Melting temperatures obtained by extrapolating ladder 5 simulations.

#### S3.3 State Probabilities and the Effect of Initial Conditions

In this section, the precision of the *probability estimates* obtained using the jackknife and bootstrap methods are provided. Additionally, the mean probability estimates are compared with the reference probabilities obtained in fig. 4 using the RMS error.

##### S3.3.1 Mean Replica Count per Metastable State (Rosta and Hummer)

The expected replica count per metastable state based on Rosta and Hummer [2] should be approximately constant with fluctuations about the mean. Since six replicas is too few to achieve the mean probabilities of states over the temperature ladders, the fluctuation amplitude and frequency are expected to be large for all states.

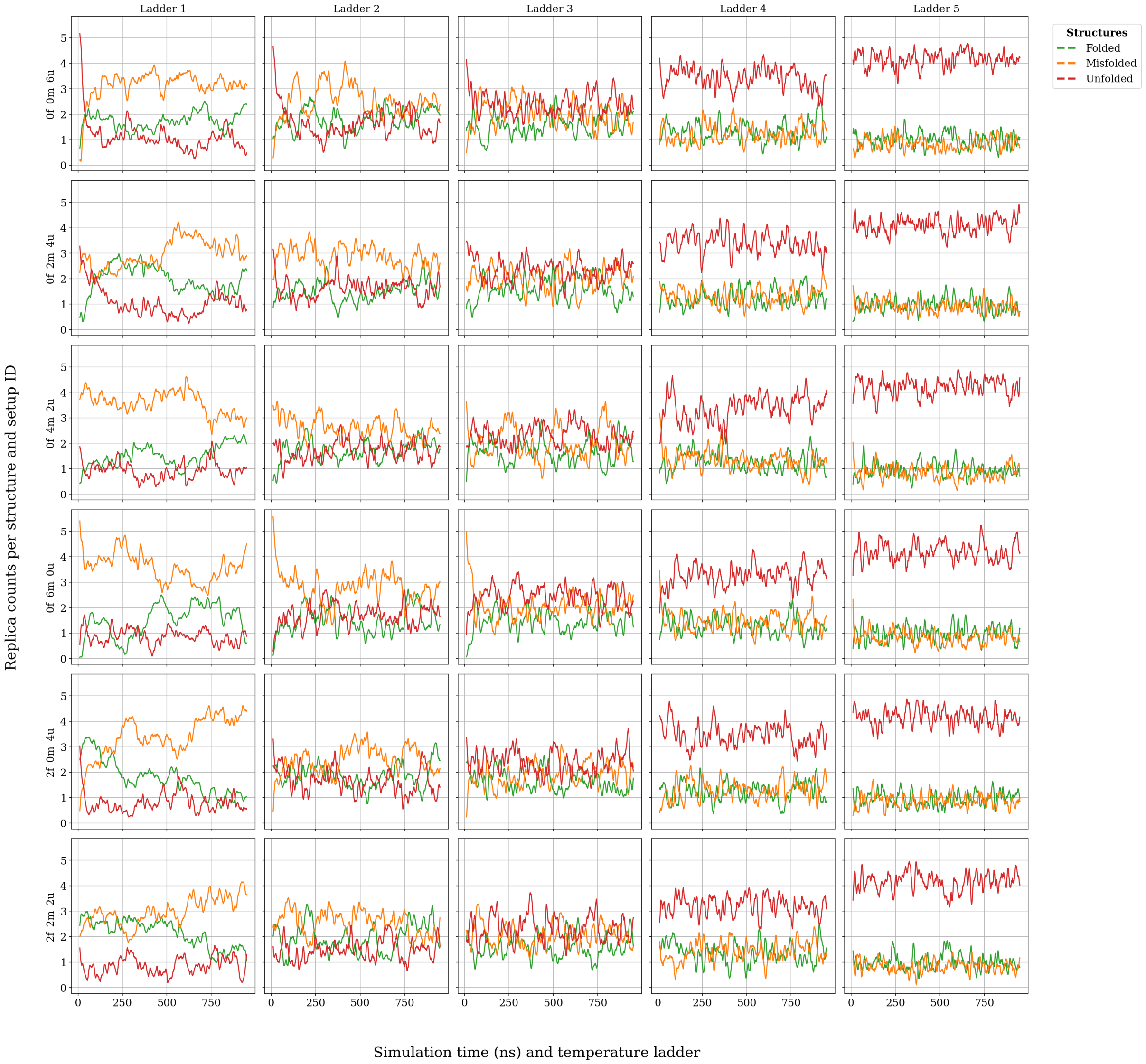

**Fig. S14:** The number of replica in each metastable state as a function of simulation time. For each setup i.e. set of starting structures and temperature ladder, the count of replicas per metastable state was calculated with time. Then, these counts were averaged over five repetitions of each setup. Finally, the running average of the "ensemble" averaged time series over a 100 frame window was calculated.

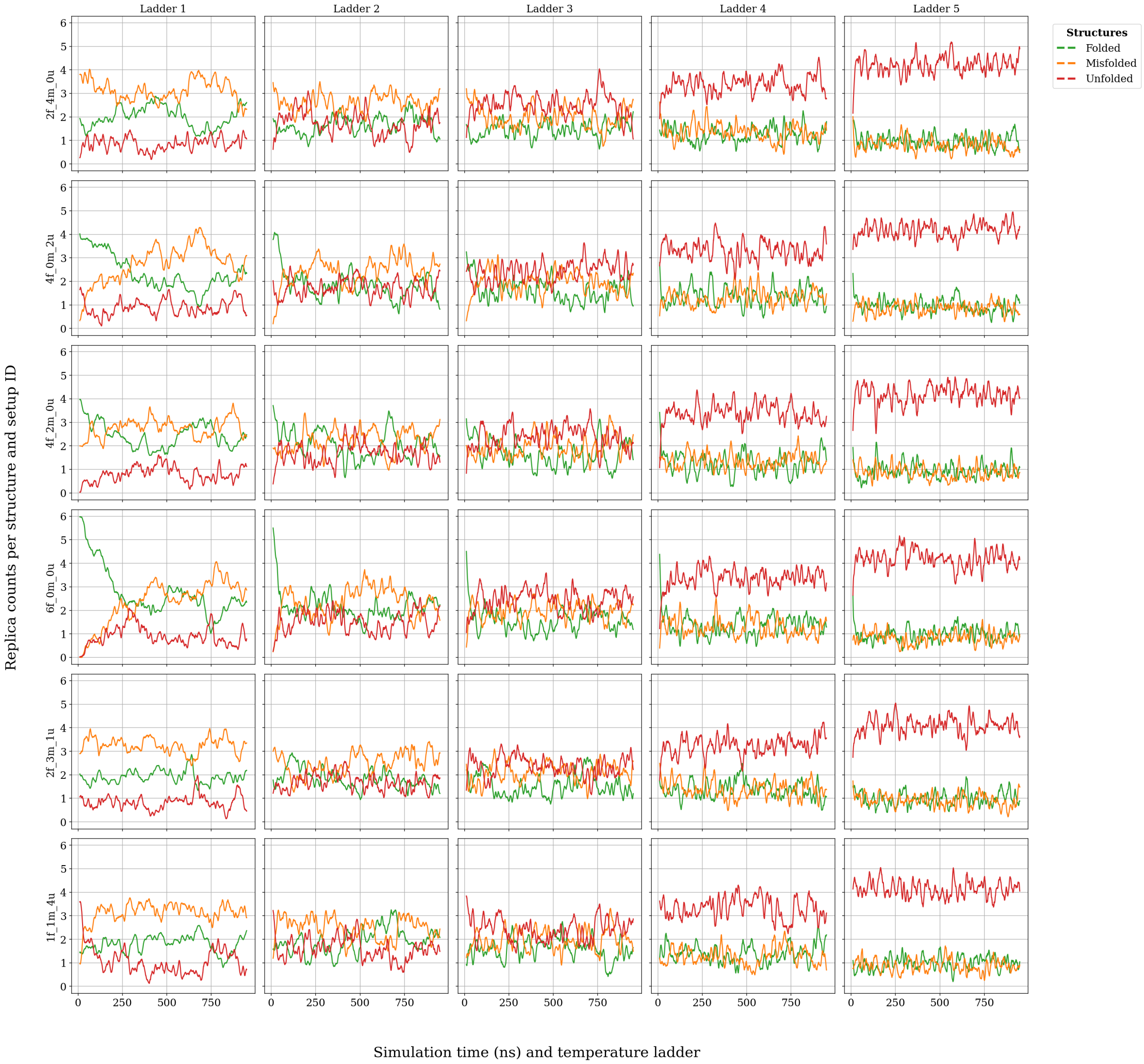

**Fig. S15:** The number of replica in each metastable state as a function of simulation time. For each setup, i.e. set of starting structures and temperature ladder, the count of replicas per metastable state was calculated with time. Then, these counts were averaged over five repetitions of each setup. Finally, the running average of the "ensemble" averaged time series over a 100 frame window was calculated.

##### S3.3.2 The Jackknife Error

The jackknife error was calculated using 20 blocks for the first ladder and 38 for the others. Calculations require at least two blocks, hence the precision curves vary from block count 2. Once estimates were made for each repetition, they were averaged.

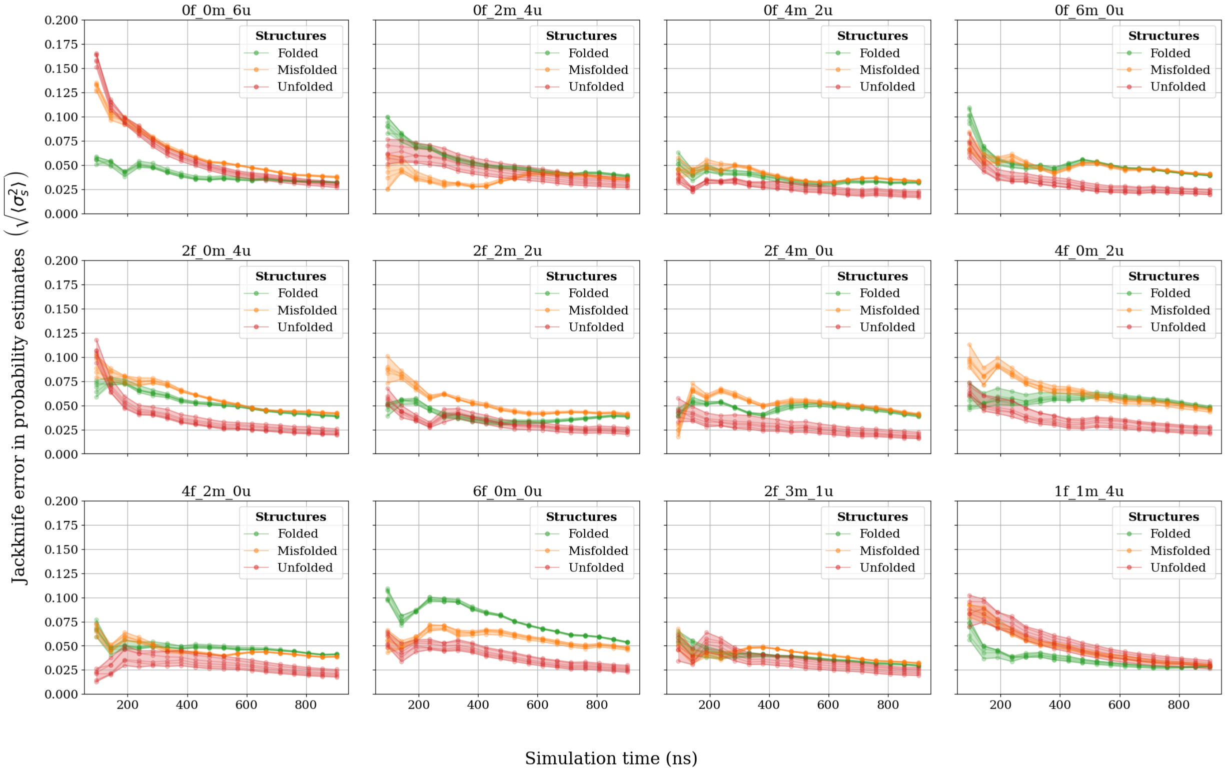

**Fig. S16:** The jackknife error for ladder 1 simulations.

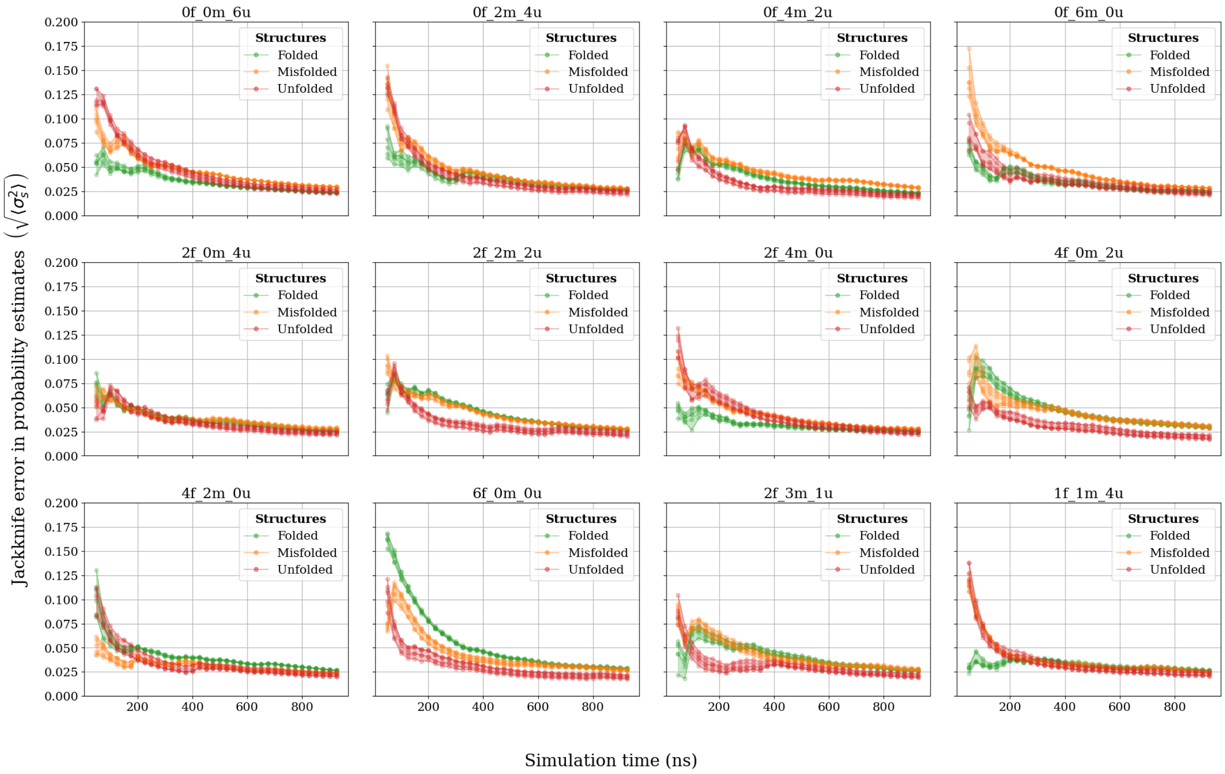

**Fig. S17:** The jackknife error for ladder 2 simulations.

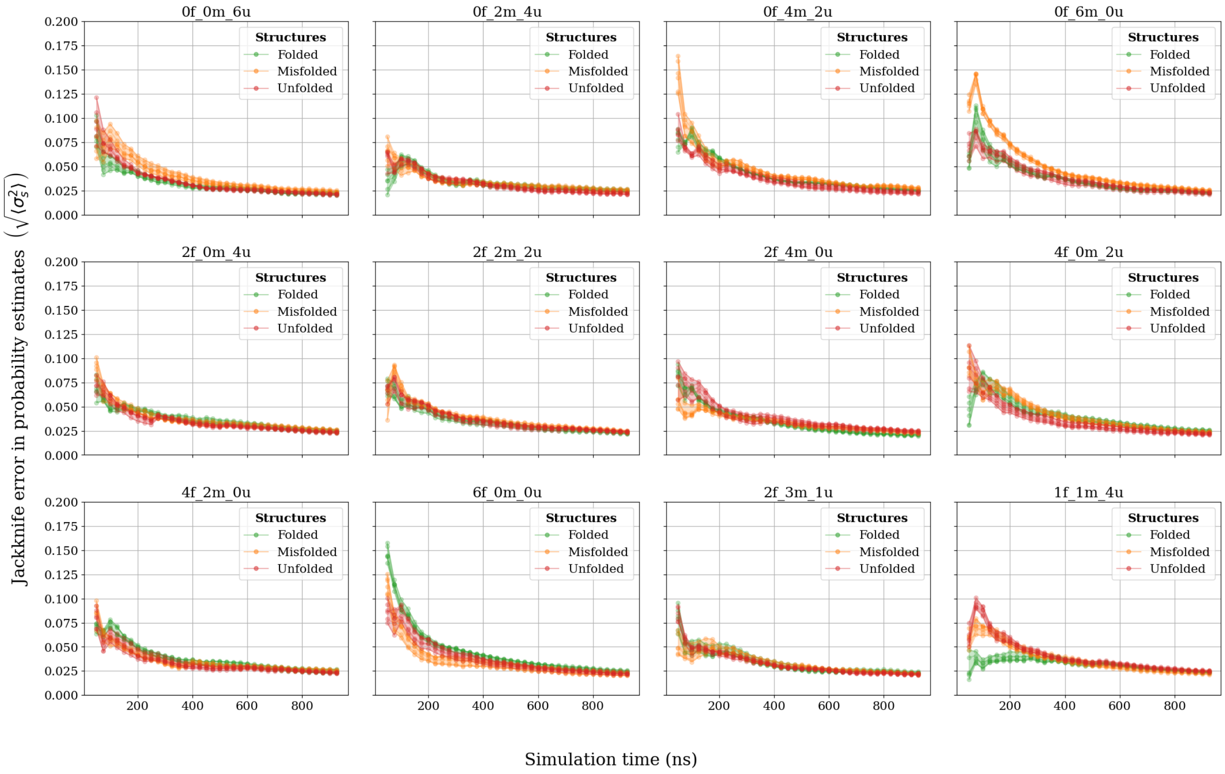

**Fig. S18:** The jackknife error for ladder 3 simulations.

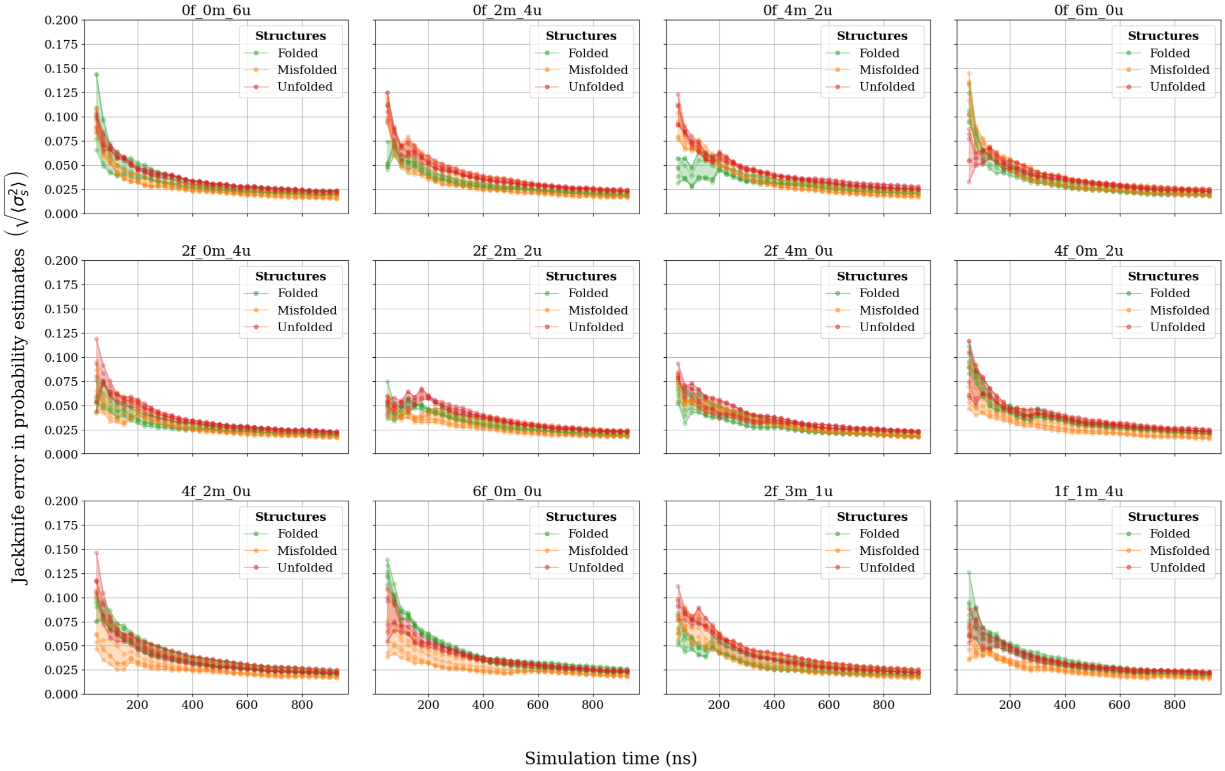

**Fig. S19:** The jackknife error for ladder 4 simulations.

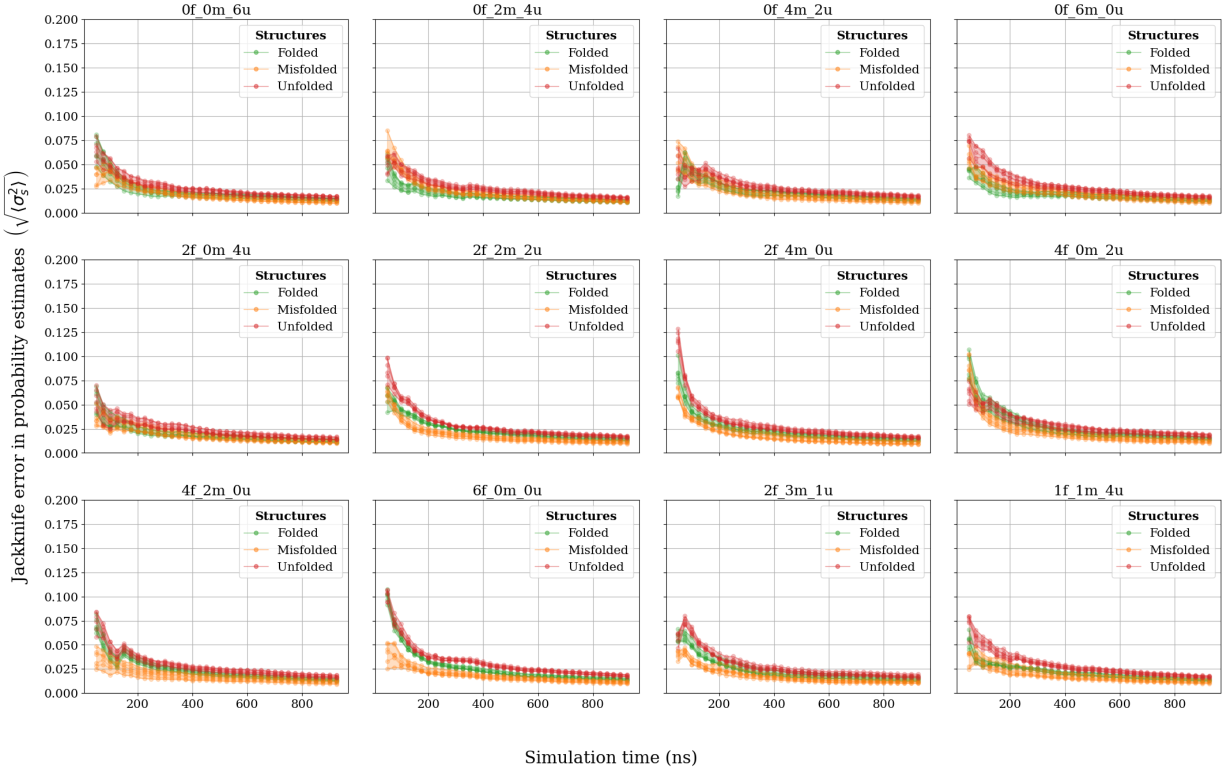

**Fig. S20:** The jackknife error for ladder 5 simulations.

##### S3.3.3 Bootstrap standard error (${}^{m}C_{m}$ errors)

The following plots show the standard error estimates for each setup (bootstrapped variance $\longrightarrow$ average over 5 repetitions $\longrightarrow$ square root). 1000 bootstraps were performed per block count.

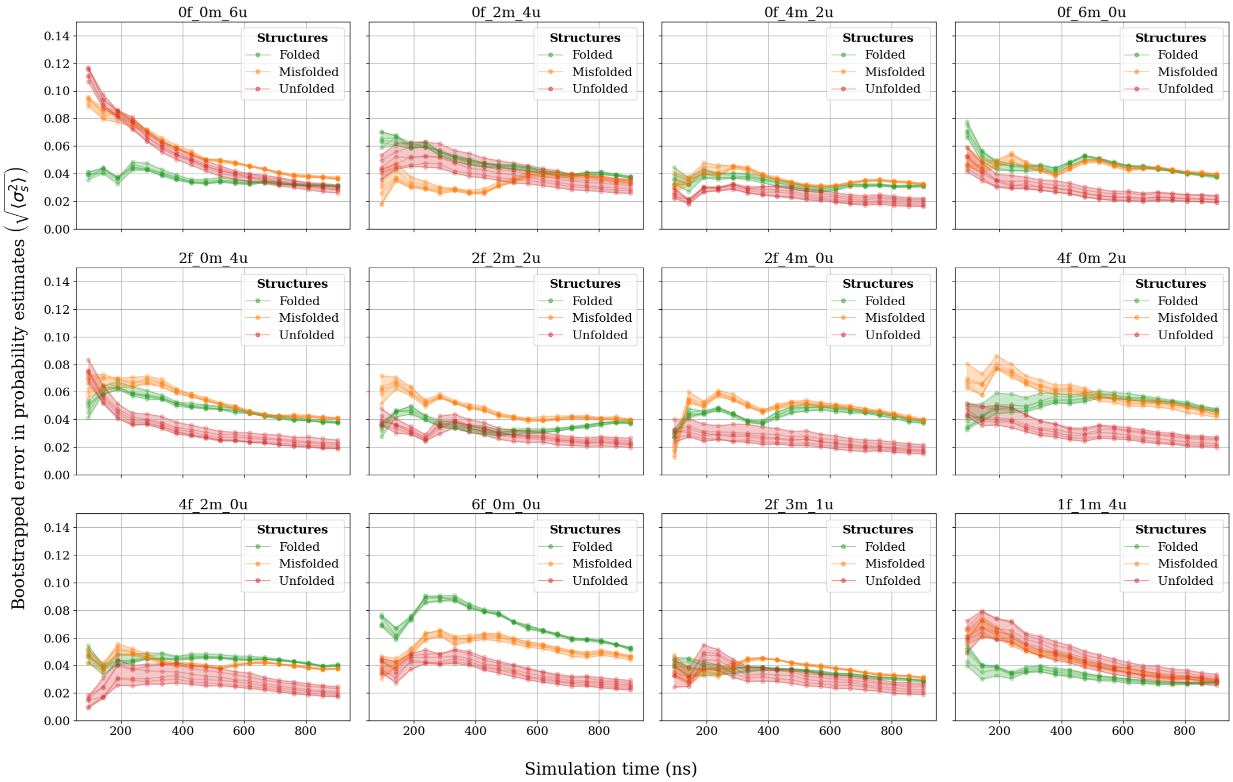

**Fig. S21:** The bootstrapped error in probability estimates for all replicas for ladder 1 simulations.

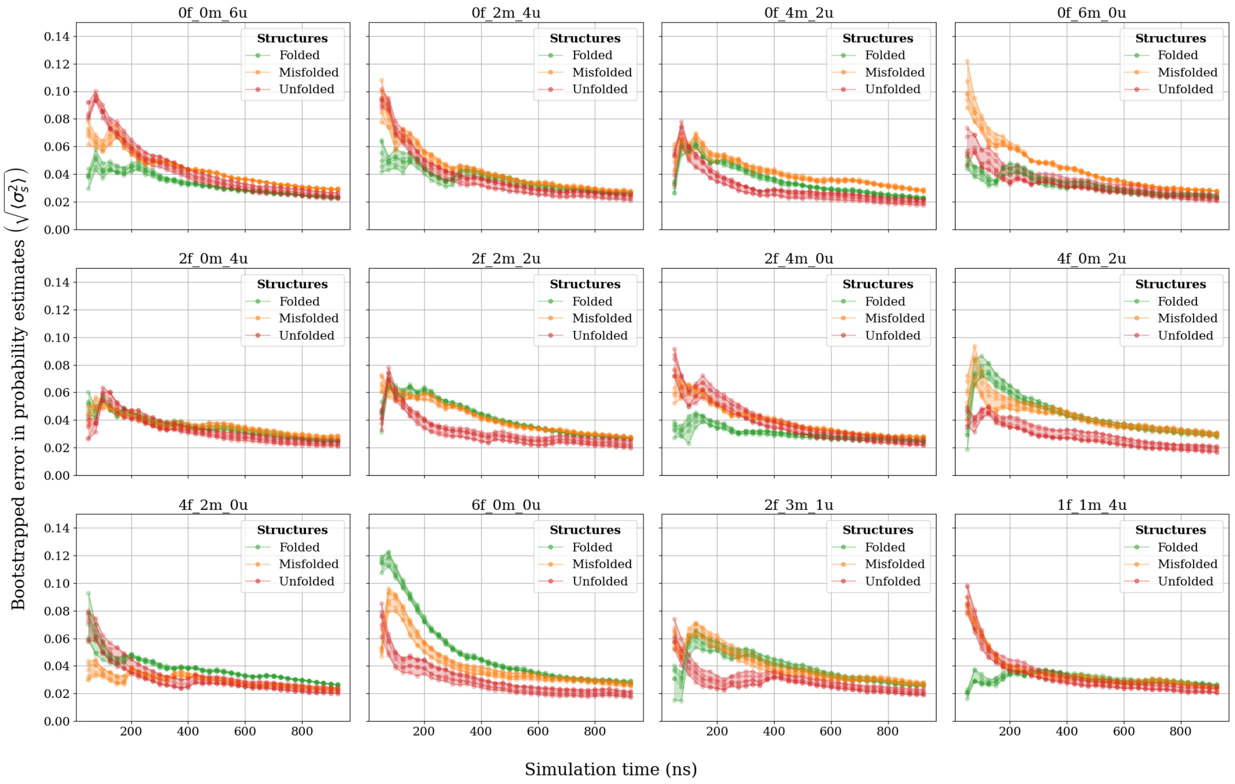

**Fig. S22:** The bootstrapped error in probability estimates for all replicas for ladder 2 simulations.

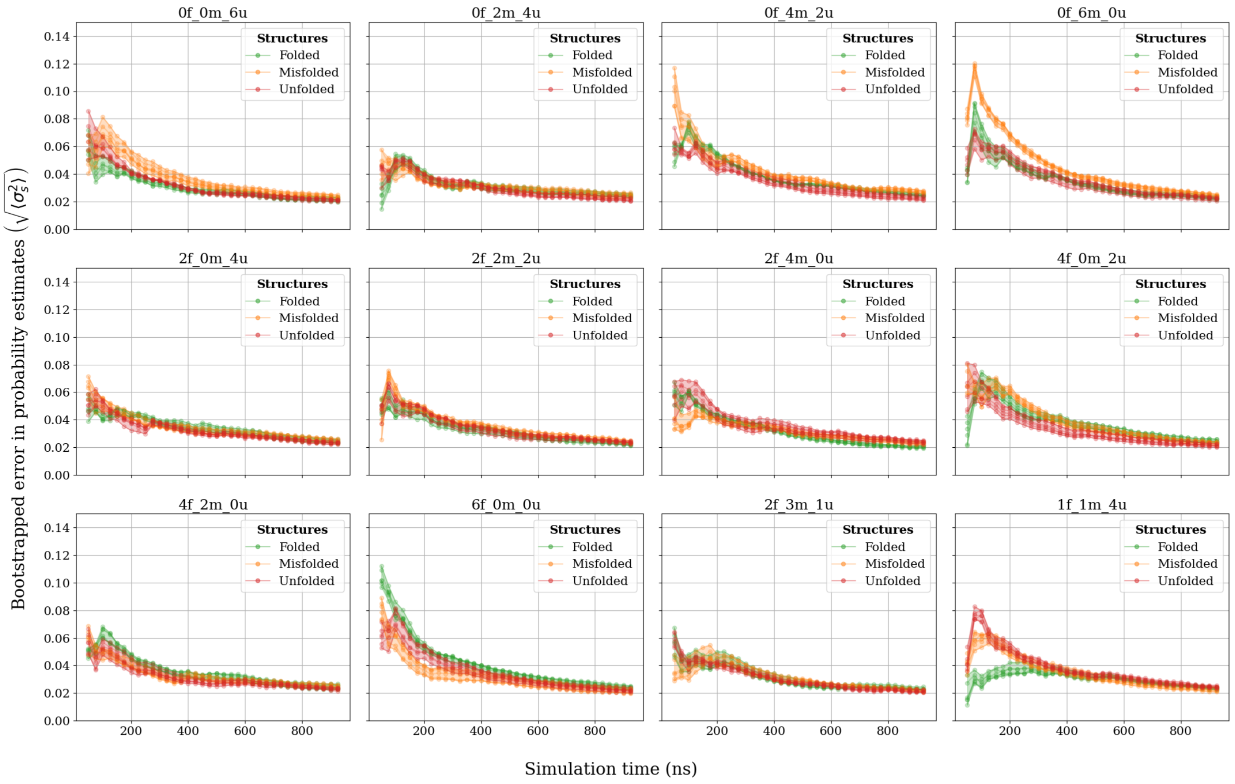

**Fig. S23:** The bootstrapped error in probability estimates for all replicas for ladder 3 simulations.

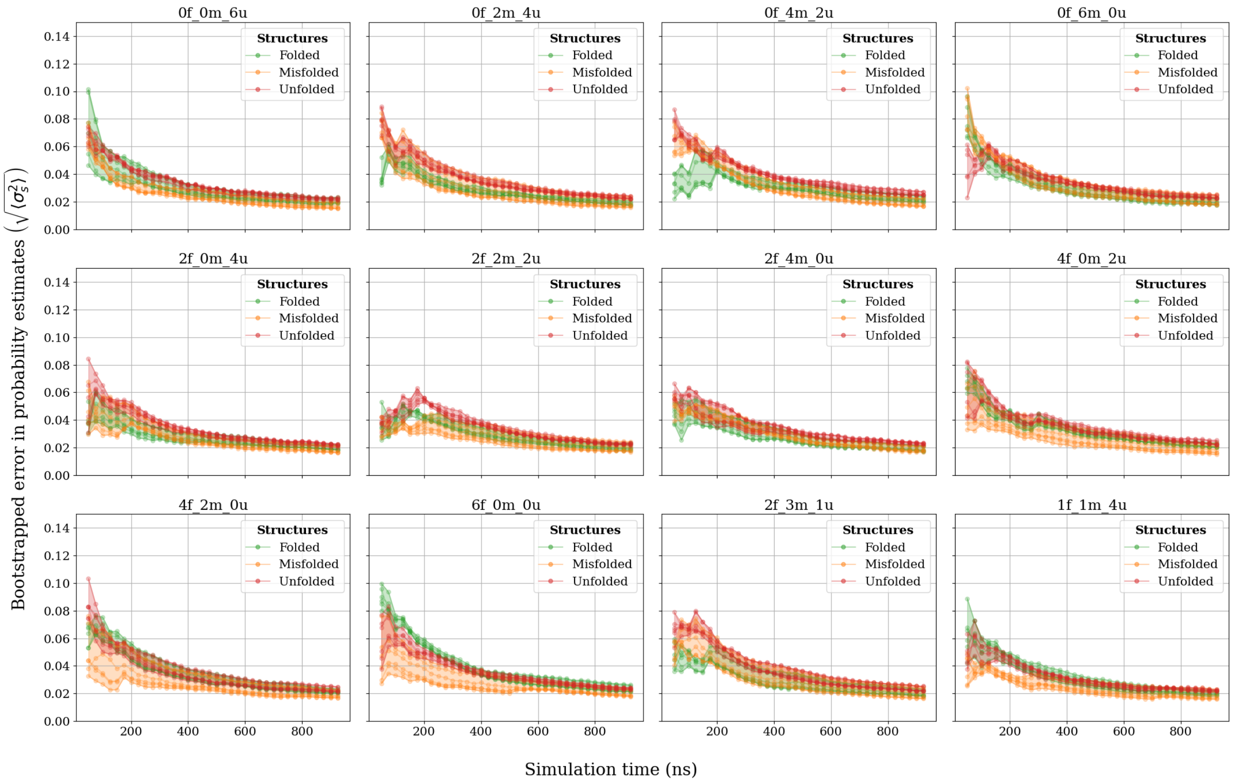

**Fig. S24:** The bootstrapped error in probability estimates for all replicas for ladder 4 simulations.

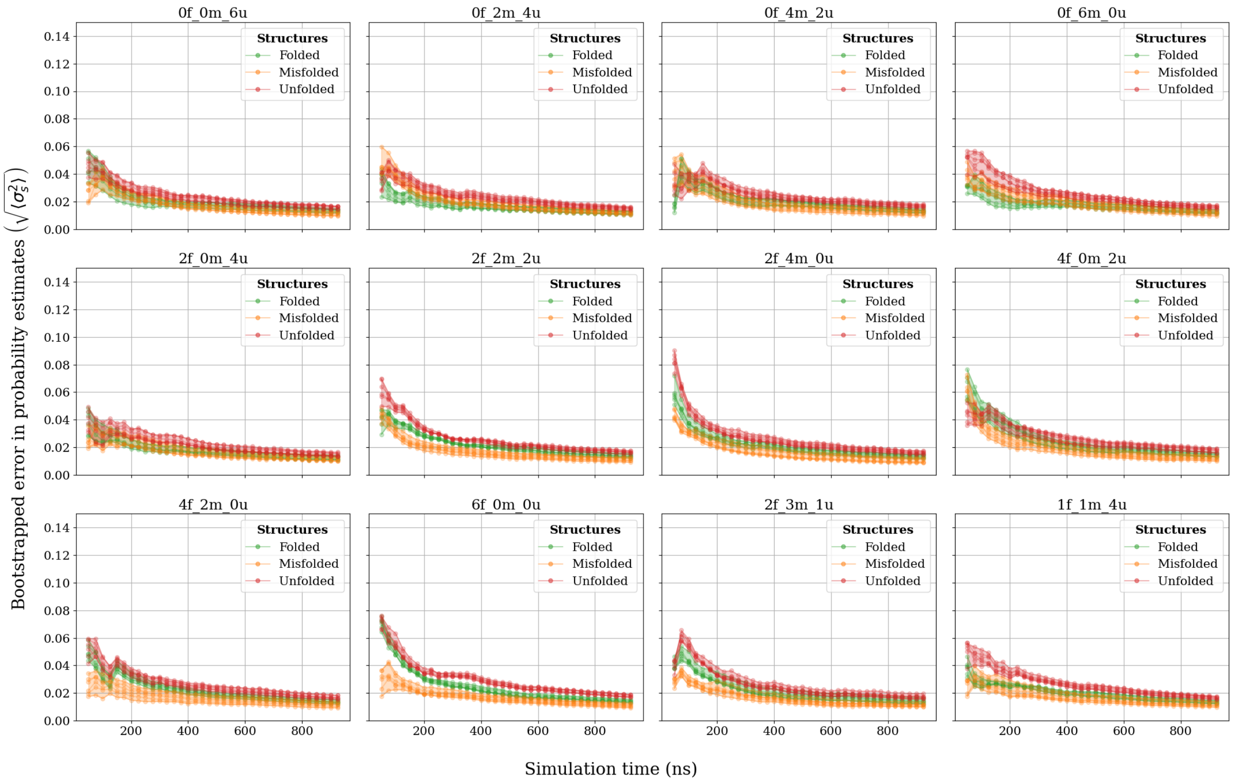

**Fig. S25:** The bootstrapped error in probability estimates for all replicas for ladder 5 simulations.

##### S3.3.4 Bootstrap RMSE

Bootstrap ⟶ mean probability for each block index ⟶ RMSE for whole ladder per block index with reference to fig. 4.

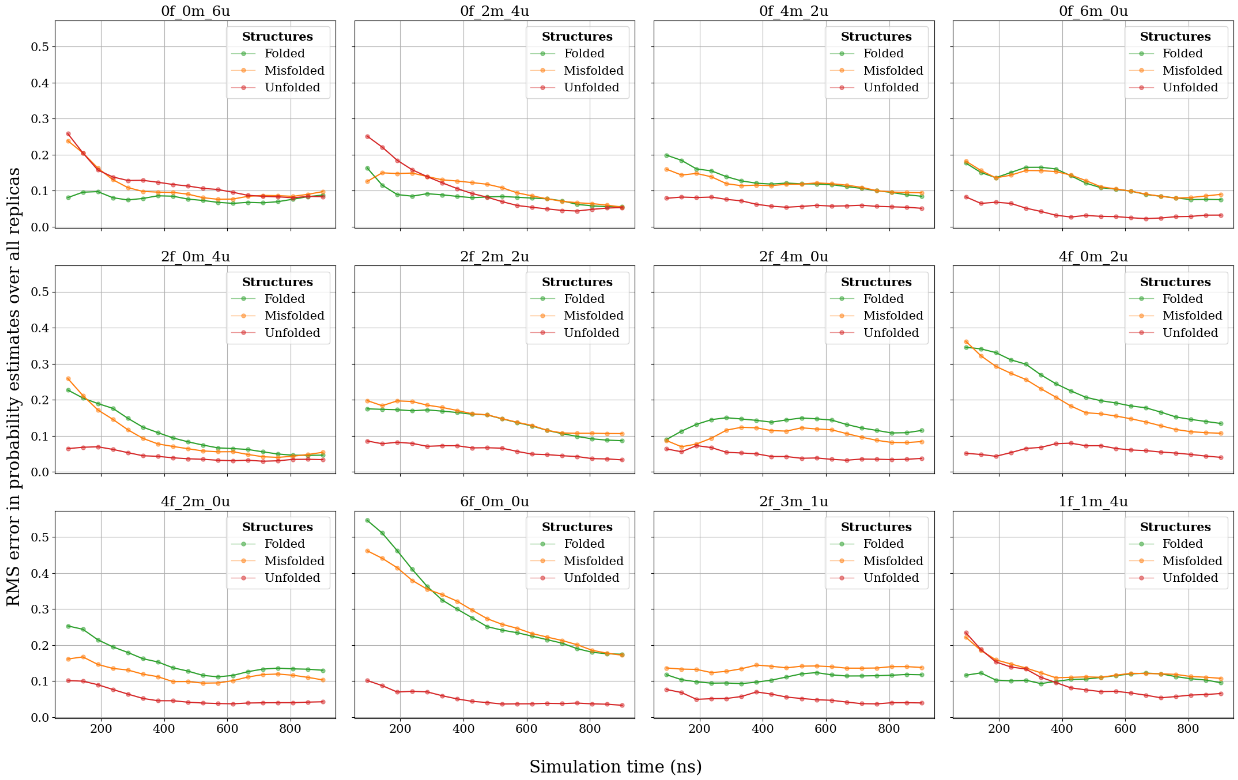

**Fig. S26:** RMSE of ladder 1 bootstrapped probability estimates with respect to reference probabilities.

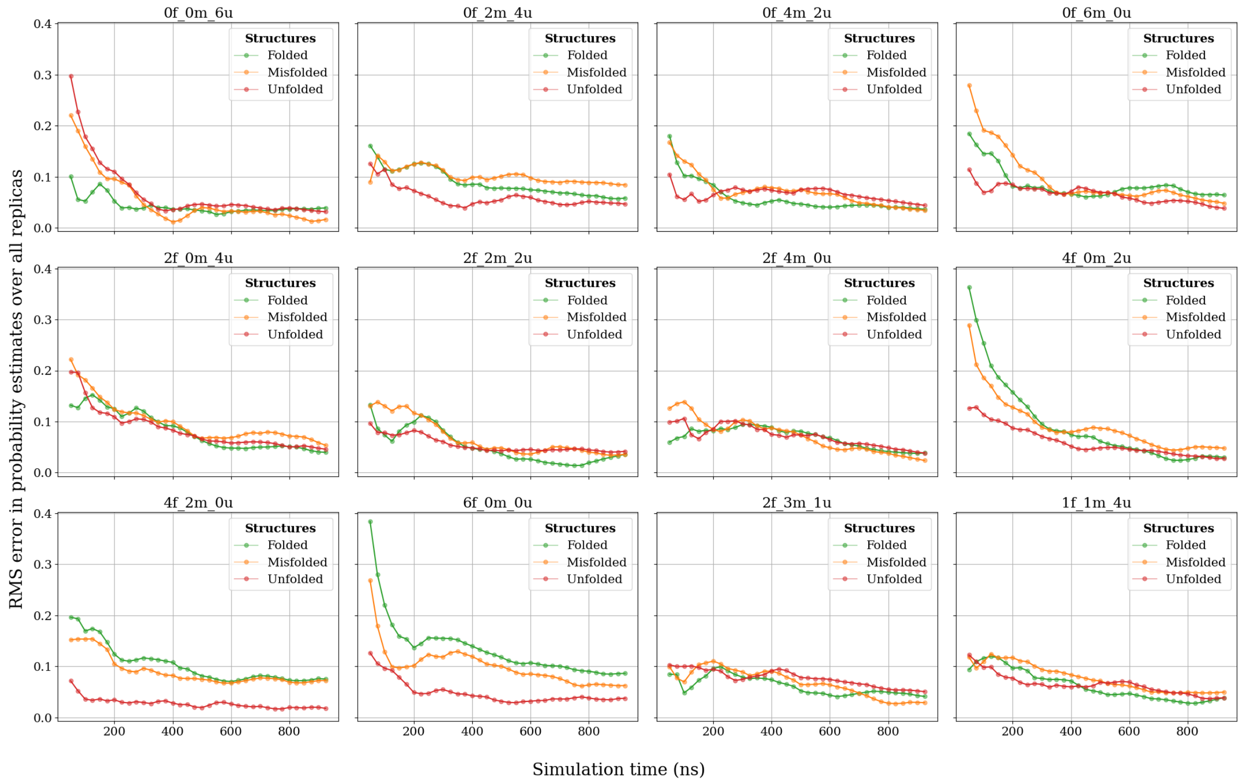

**Fig. S27:** RMSE of ladder 2 bootstrapped probability estimates with respect to reference probabilities.

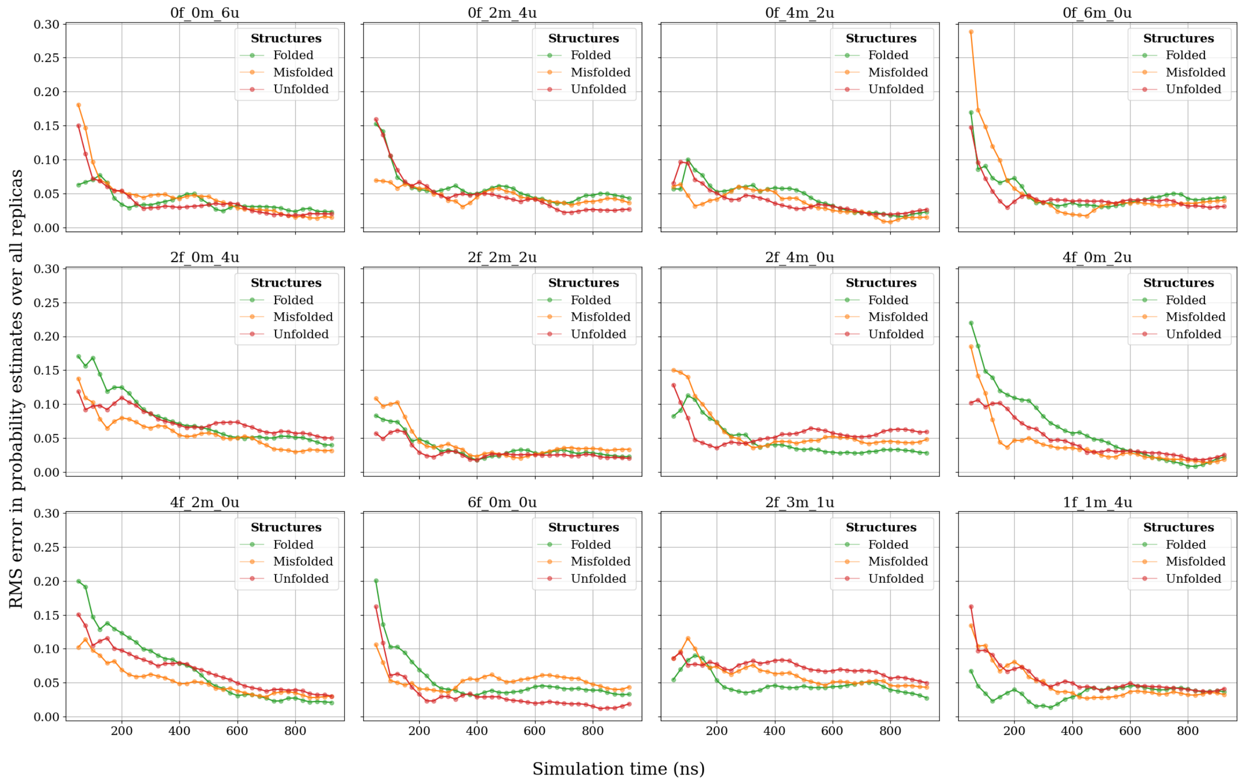

**Fig. S28:** RMSE of ladder 3 bootstrapped probability estimates with respect to reference probabilities.

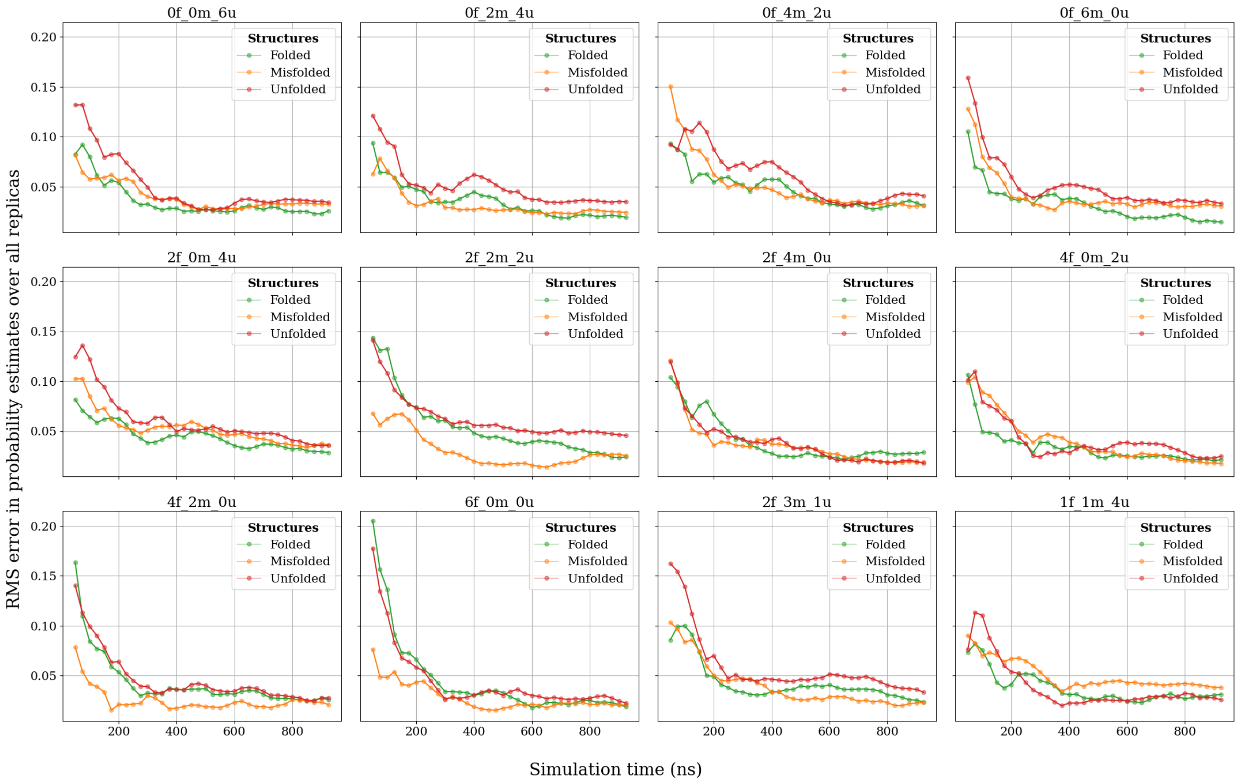

**Fig. S29:** RMSE of ladder 4 bootstrapped probability estimates with respect to reference probabilities.

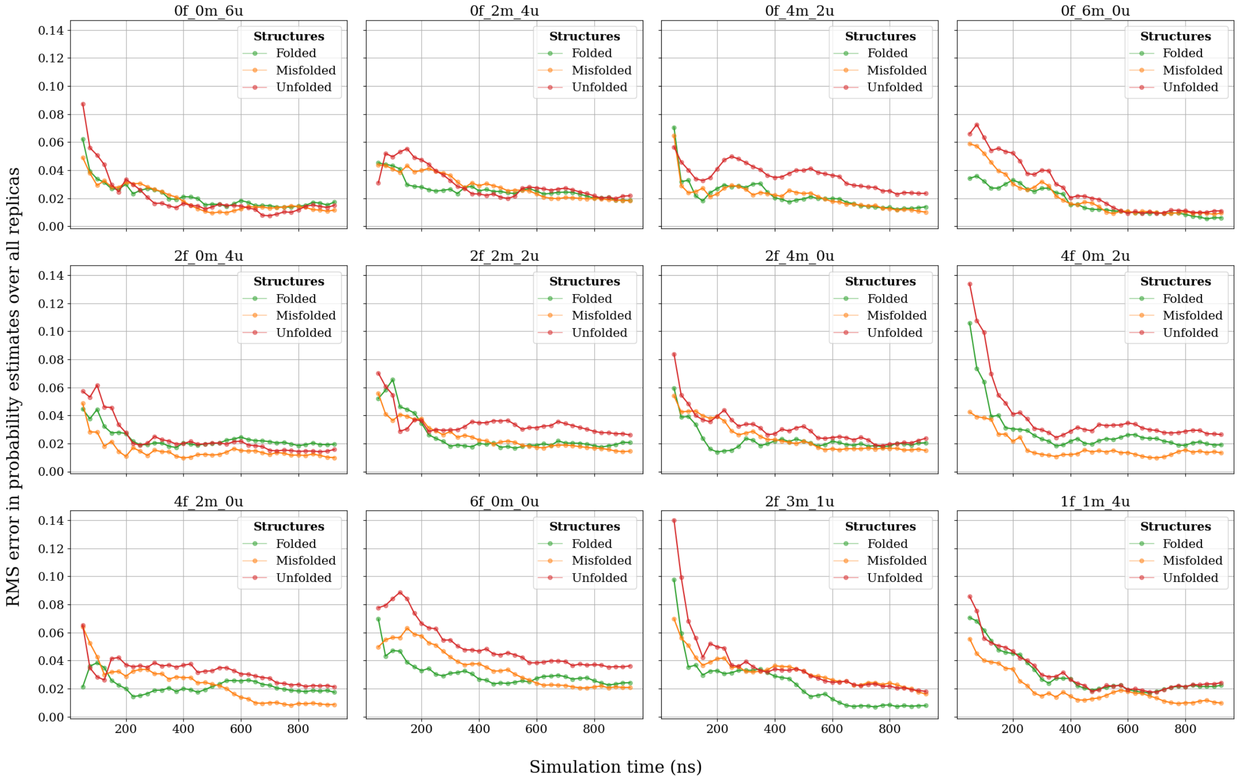

**Fig. S30:** RMSE of ladder 5 bootstrapped probability estimates with respect to reference probabilities.

##### S3.4 Supplementary Materials for Chignolin Simulations with FF14SB and FF19SB

The $T_M$ estimates obtained by combining the two temperature ladders has been provided in fig. S32. It can be seen that the uncertainty of the estimates are fairly stable. Despite overestimating the fraction of the folded and misfolded states in both ladders, it is observed that the $T_M$ values actually appear converged.

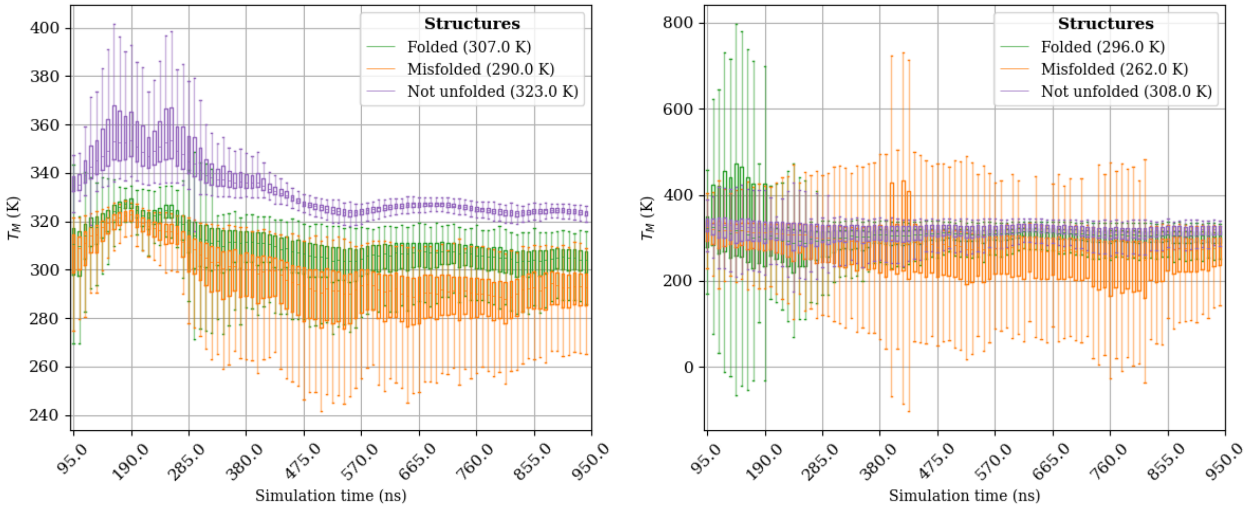

**(a)** Ladder 1

**(b)** Ladder 2

**Fig. S31:** The $T_M$ estimates obtained from the van't Hoff analysis of the different temperature ladders with FF14SB. The estimates obtained from the high temperature ladder are inaccurate and imprecise.

The final estimates are approximately 303±2 K, 294±3 K and 323±1 K for the folded, misfolded and not-unfolded states respectively. The estimates, particularly those of the (natively) folded and not unfolded states are close to the experimental $T_M$ of 310 K-315 K found in Honda et al. [3].

The $T_M$ estimates obtained from the low temperature and high temperature ladders taken individually with the FF19SB force field have been provided in fig. S33. It is clear that the estimates are very uncertain and that large fluctuations are observed at all points in the simulation. The combined estimate is provided in fig. S34 and it is once more clear that the results are unreliable. This is a common problem that has been discussed in section 4.3.

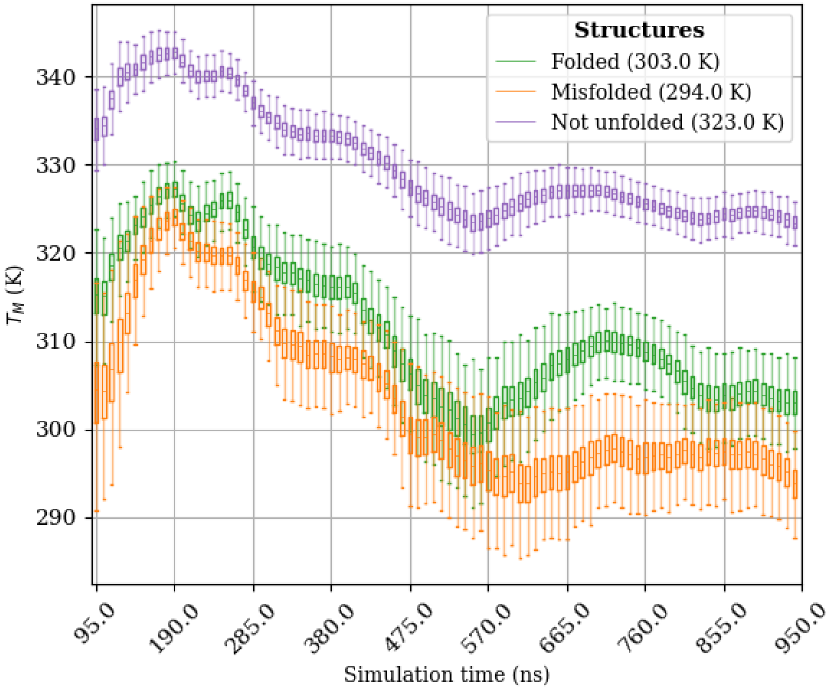

**Fig. S32:** $T_M$ estimates of Chignolin obtained by combining both ladders with the FF14SB force field. The overall uncertainty is lower than for the individual ladders.

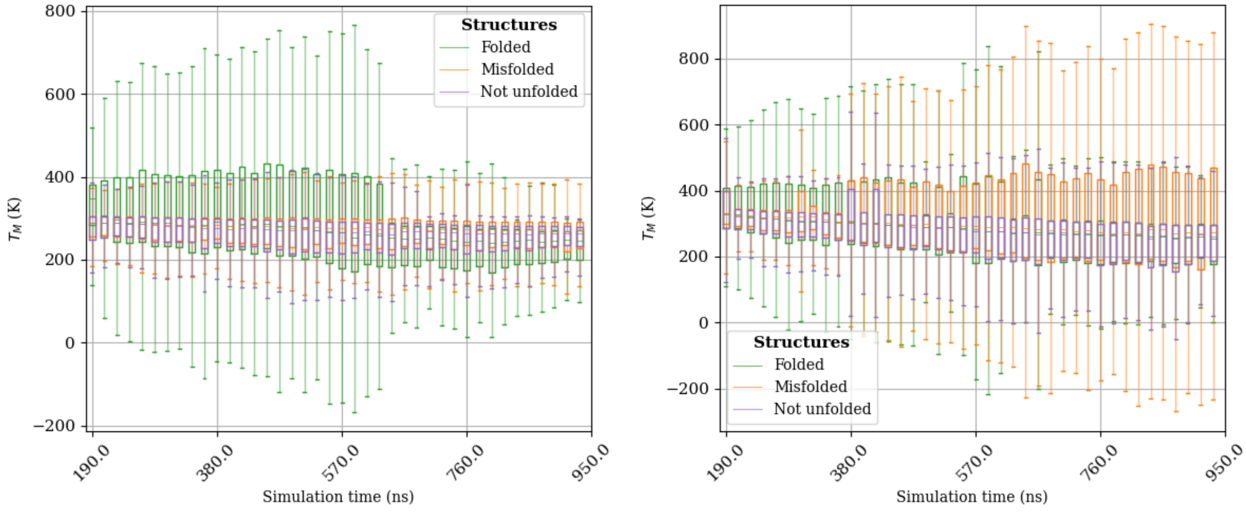

**(a)** Ladder 1

**(b)** Ladder 2

**Fig. S33:** The melting temperature estimates with the FF19SB force field using single ladders. The estimates are highly uncertain and unphysical.

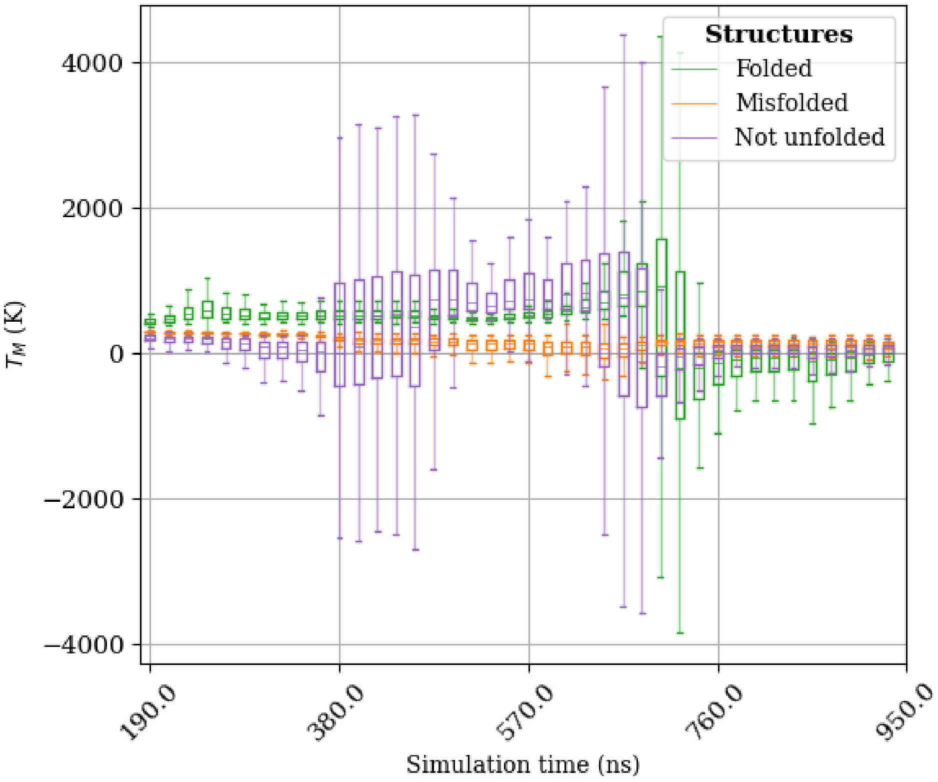

**Fig. S34:** $T_M$ estimates of Chignolin obtained by combining both ladders with the FF19SB force field.

[2] Edina Rosta and Gerhard Hummer. Error and efficiency of replica exchange molecular dynamics simulations. 131(16):165102. ISSN 0021-9606. doi: 10.1063/1.3249608.

[3] Shinya Honda, Kazuhiko Yamasaki, Yoshito Sawada, and Hisayuki Morii. 10 Residue Folded Peptide Designed by Segment Statistics. 12(8):1507–1518. ISSN 09692126. doi: 10.1016/j.str.2004.05.022.
